# The autism-associated gene *Scn2a* plays an essential role in synaptic stability and learning

**DOI:** 10.1101/366781

**Authors:** Perry WE Spratt, Roy Ben-Shalom, Caroline M Keeshen, Kenneth J Burke, Rebecca L Clarkson, Stephan J Sanders, Kevin J Bender

## Abstract

Autism spectrum disorder (ASD) is strongly associated with *de novo* gene mutations. One of the most commonly affected genes is *SCN2A*. ASD-associated *SCN2A* mutations impair the encoded protein Na_V_1.2, a sodium channel important for action potential initiation and propagation in developing excitatory cortical neurons. The link between an axonal sodium channel and ASD, a disorder typically attributed to synaptic or transcriptional dysfunction, is unclear. Here, we show Na_V_1.2 is unexpectedly critical for dendritic excitability and synaptic function in mature pyramidal neurons, in addition to regulating early developmental axonal excitability. Na_V_1.2 loss reduced action potential backpropagation into dendrites, impairing synaptic plasticity and synaptic stability, even when Na_V_1.2 expression was disrupted late in development. Furthermore, we identified behavioral impairments in learning and sociability, paralleling observations in children with *SCN2A* loss. These results reveal a novel dendritic function for Na_V_1.2, providing insight into cellular mechanisms likely underlying circuit and behavioral dysfunction in ASD.

## Introduction

High-throughput sequencing has implicated numerous genes in autism spectrum disorder (ASD) (Sanders et al., 2015). These genes cluster into two groups: chromatin modifiers and genes that support synaptic function (De Rubeis et al., 2014; Sanders et al., 2015). Surprisingly, *SCN2A*, the gene with the most robust ASD association (Ben-Shalom et al., 2017; De Rubeis et al., 2014; Sanders et al., 2015), does not readily fit within either group. *SCN2A* encodes the protein Na_V_1.2, a voltage-gated sodium channel with a known role in the axon initial segment (AIS), an axonal subcompartment adjacent to the soma that is the site of action potential (AP) initiation (Bender and Trussell, 2012). Interestingly, *SCN2A* variants with different effects on channel function are associated with distinct neurodevelopmental disorders. Heterozygous loss of function mutations (haploinsufficiency) in *SCN2A* that diminish or eliminate channel function are strongly associated with ASD as well as intellectual disability (ID), whereas gain of function missense mutations are strongly associated with infantile epilepsy of varying severity (Ben-Shalom et al., 2017; Sanders et al., 2018; Wolff et al., 2017). While neuronal hyperexcitability due to gain-of-function *SCN2A* variants likely contributes to infantile epilepsy, the neuropathological mechanisms underlying the strong association between *SCN2A* loss of function and ASD/ID remains largely unknown.

In neocortex, Na_V_1.2 is primarily expressed in glutamatergic pyramidal cells (Hu et al., 2009; Li et al., 2014; Yamagata et al., 2017), where its distribution within different axonal compartments changes over development. In the first postnatal week in mice, corresponding to late gestation through the first year of life in humans (Workman et al., 2013), Na_V_1.2 is the only sodium channel isoform expressed in the axon and AIS, and is thus solely responsible for the initiation and propagation of action potentials (Gazina et al., 2015). Later in development, Na_V_1.2 in the axon and distal AIS is replaced by Na_V_1.6 (*SCN8A*), which has a lower voltage threshold for activation (Bender and Trussell, 2012; Kole and Stuart, 2012). Consequently, the distal AIS becomes the site of AP initiation and Na_V_1.2, now restricted to the proximal AIS, is thought to promote effective backpropagation of APs from the AIS into the soma (Hu et al., 2009). Given these potentially distinct developmental roles, it is critical to understand how *SCN2A* haploinsufficiency affects neuronal excitability and network function across development, as this may shed light on ASD and ID etiology.

Here, we examined cellular, synaptic, and behavioral consequences of heterozygous loss of *Scn2a* in mice (*Scn2a^+/-^*). AP initiation was impaired in early development, consistent with a major role for *Scn2a* in axonal excitability. Unexpectedly, we identified deficits in somatodendritic excitability, consistent with the expression of Na_V_1.2 channels throughout these compartments, that arose after early development and persisted throughout life. Furthermore, excitatory synapses onto *Scn2a^+/-^* neurons were both functionally and structurally immature in adulthood. Similar synaptic impairments were found even when *Scn2a* expression was disrupted late in development, selectively impairing dendritic excitability without a period of reduced axonal excitability. Reduced dendritic excitability impaired the backpropagation of action potentials into distal dendrites, with corresponding deficits in spike-timing dependent plasticity. Finally, we observed sex-specific deficits in both learning and sociability in *Scn2a^+/-^* mice. Together, these results identify a novel role for Na_V_1.2 in the dendrites of mature pyramidal neurons, demonstrating that Na_V_1.2 contributions to dendritic, and not only axonal, excitability is essential for proper synaptic development and function. This generates the hypothesis that *SCN2A* contributes to ASD and ID by disrupting synaptic function, like many other ASD genes, and that these effects persist in mature neurons.

## Results

### *Scn2a* loss impairs axonal and dendritic excitability in distinct developmental periods

To determine how *Scn2a* haploinsufficiency affects cortical neuron excitability, acute slices containing medial prefrontal cortex were prepared from *Scn2a^+/-^* and wild-type (WT) littermate control mice aged postnatal day (P)4 through P64. *Scn2a^+/-^* mice have a 50% reduction in Na_V_1.2 mRNA, have reduced Na_V_-mediated currents in dissociated cell culture at P5-9 (Planells-Cases et al., 2000), and are seizure free (Mishra et al., 2017), consistent with the majority of *Scn2a* loss-of-function cases reported in ASD (Sanders et al., 2018). We focused our studies primarily on subcortically projecting layer 5b thick-tufted neurons that were identified based on electrophysiological properties (see Methods), as dysfunction within this region and cell class has been implicated in ASD (Willsey et al., 2013). Neuronal excitability was assessed using a series of current steps to generate APs. AP threshold was depolarized and spike output was reduced relative to WT in the first postnatal week (Fig. 1A, C, F). Differences between *Scn2a^+/-^* and WT were not present thereafter, consistent with increased Na_V_1.6 expression in the AIS. In agreement, persistent sodium currents, which reflect the recruitment of sodium channels with the lowest threshold (Taddese and Bean, 2002), were suppressed in *Scn2a^+/-^* mice at P6, and not P30 (Fig. S1A-D). Furthermore, axonal conduction, assayed by imaging AP-evoked calcium transients in boutons, was reliable in mature neurons (Fig. S2). Input resistance was comparable between *Scn2a^+/-^* and WT at all ages, and transient and sustained potassium currents were no different in mature neurons (Fig. S1E-G). Overall, these data suggest that *Scn2a* haploinsufficiency impairs neuronal excitability early in development without functional compensation from the remaining *Scn2a* allele or other ion channels involved in AP initiation.

**Figure 1:**
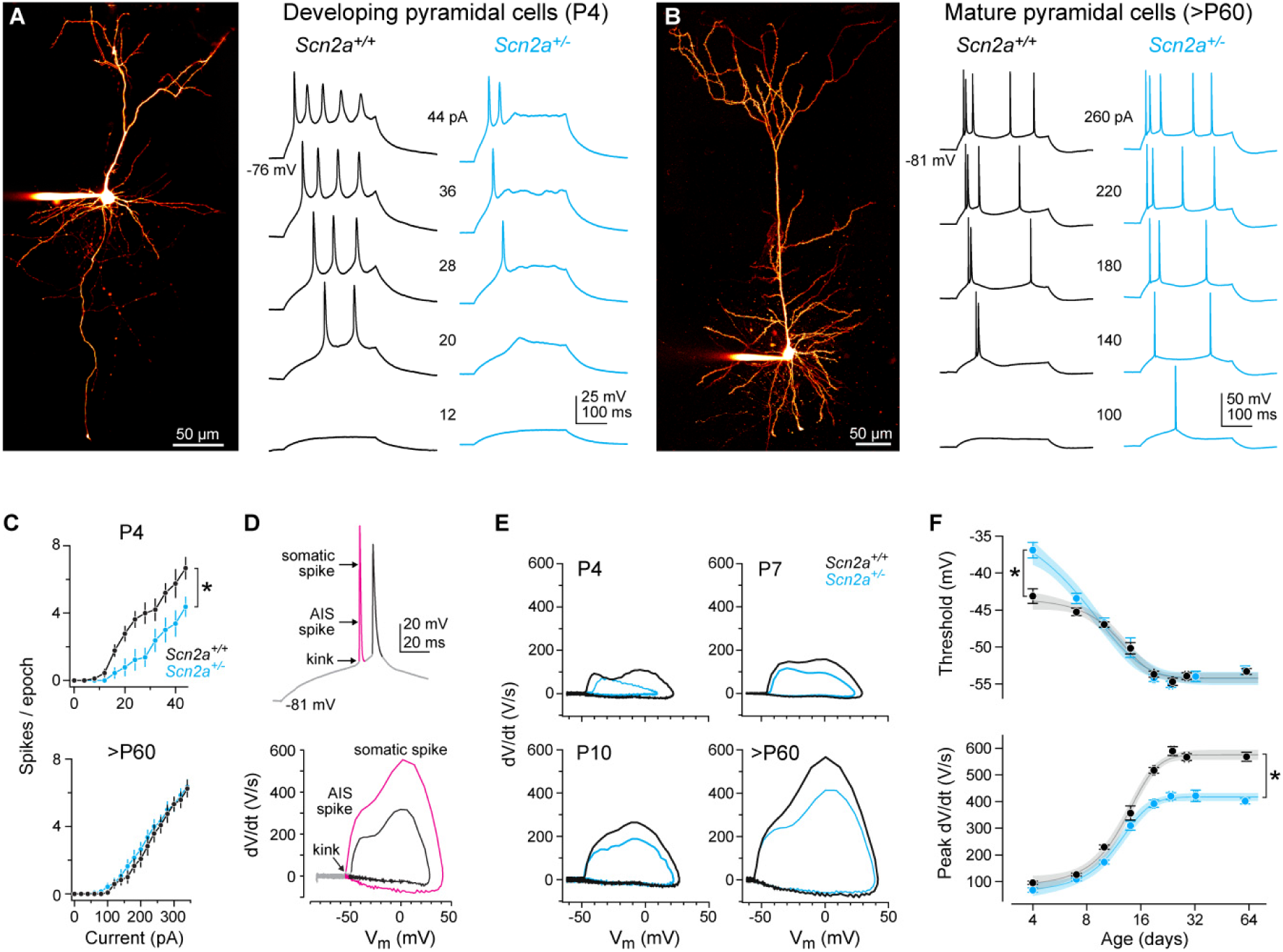
*Scn2a* haploinsufficiency impairs different aspects of neuronal excitability across developmental periods. **A:** Left, 2-photon laser scanning microscopy (2PLSM) z-stack of developing pyramidal cell (P7). Right, APs generated by current injection (12–44 pA, 300 ms) in *Scn2a^+/+^* (black) and *Scn2a^+/-^* (cyan) cells. AP data from neurons recorded at P4. **B:** Left, 2-photon z-stack of mature thick-tufted pyramidal cell. Right, AP response as in (A). **C:** APs (spikes) per 300 ms stimulation epoch for each current amplitude. At P4, *Scn2a^+/-^* are less excitable (Firing rate slope between 4–44 pA, *Scn2a^+/+^*: 0.67 ± 0.07 APs/pA*s, n = 9; *Scn2a^+/-^*: 0.41 ± 0.07, n = 9, *: p = 0.02, Two-sided Mann-Whitney). At >P60, no differences are observed between *Scn2a^+/+^* and *Scn2a^+/-^* cells (Slope between 50–350 pA: *Scn2a^+/+^*: 0.08 ± 0.007 APs/pA*s, n = 12; *Scn2a^+/-^*: 0.08 ± 0.005, n = 14, p = 0.96, Two-sided Mann-Whitney). **D:** A burst of 2 APs plotted as voltage vs. time (top) and dV/dt vs voltage (phase-plane plot, bottom). Each AP is color-coded across panels, and aspects of rising phase of the AP corresponding to initiation of AP in AIS and full detonation of AP in soma are highlighted. **E:** Phase-plane plots at P4, P7, P10, and >P60 in *Scn2a^+/+^* and *Scn2a^+/-^* cells. Note recovery of AP threshold (kink) deficit between P4 and P10, and decrements in peak dV/dt that become more pronounced with age. **F:** Top: AP threshold across development from P4-64 (log2 scale for age) in *Scn2a^+/+^* (black) and *Scn2a^+/-^* (cyan) mice. Bottom: Peak rising phase dV/dt vs. age. Circles and bars are mean ± SEM within an age group (n = 7-28 cells per group). Data fit with sigmoid function with 95% confidence intervals determined from fits to single cells. *: difference in parameters of the sigmoid fits. Top: Left asymptote, *Scn2a^+/+^*: -44.1 ± 0.6 mV, *Scn2a^+/-^* -35.6 ± 1.6, p < 0.001, unpaired two-sided t-test. Bottom: Right asymptote, *Scn2a^+/+^*: 587.8 ± 10.2 V/s, *Scn2a^+/-^*: 420.3 ± 9.5, p < 0.001, unpaired two-sided t-test.

Although impairments in AP threshold recovered after P7, a closer examination of the AP waveform revealed a striking reduction in the velocity of the AP (dV/dt) that became more pronounced as neurons matured (Fig. 1E-F) (>P22, WT: 578.2 ± 9.2 V/s, n = 44; *Scn2a^+/-^*: 417 ± 9.5, n = 33; p < 0.001, Two-sided Mann-Whitney). Similar relative deficits were identified in putative thin-tufted neurons, which have different AP waveform characteristics compared to thick-tufted neurons (Clarkson et al., 2017) (>P18: WT: 426.9 ± 24.7 V/s, n = 8; *Scn2a^+/-^*: 358 ± 21.1, n = 9; p = 0.007, Two-sided Mann-Whitney). Plotting AP velocity as a function of voltage (phase-plane) highlights two components of the rising phase of the AP, as measured with somatic current-clamp. The first relates to the initiation of the AP in the AIS and the second to the recruitment of somatic sodium channels (Bean, 2007). In *Scn2a^+/-^* cells, both AIS and somatodendritic components of the AP were slower, suggesting that Na_V_1.2 channels are distributed in both the AIS and somatodendritic compartment, and that their density is lower in both compartments. Similar deficits were not observed in parvalbumin or somatostatin positive interneurons (Fig. S3), consistent with a lack of Na_V_1.2 expression in these cell classes (Yamagata et al., 2017; but see Li et al., 2014). Thus, in mature PFC networks, *Scn2a* haploinsufficiency may be more consequential for somatodendritic excitability than axonal excitability in excitatory neurons.

To explore the cellular consequences of reduced somatodendritic excitability, we modified established computational models of cortical pyramidal cells by varying the localization and relative density of Na_V_1.2 and Na_V_1.6 (Ben-Shalom et al., 2017). First, Na_V_1.2 channels were localized only to the AIS, and AP waveform was compared between models with 100% and 50% Na_V_1.2 expression. AP waveform was only modestly altered by such manipulations. To best account for empirical observations, we found that Na_V_1.2 expression was required throughout the somatodendritic compartment, in equal levels with Na_V_1.6 (Fig. 2A-B, Fig. S4A-E). In this configuration, we modeled AP waveform throughout the dendritic arbor and found that backpropagating APs (bAPs) from the AIS to the dendritic arbor were attenuated in a distance dependent fashion. Though AP amplitudes were reduced only 10% in the soma, they were attenuated to far greater extents in distal dendrites (41%). These bAPs provide an instructive signal for dendritic integration and synaptic plasticity, largely through their ability to recruit dendritic voltage-dependent calcium channels and NMDA receptors (Feldman, 2012; Larkum et al., 1999). Since this recruitment is nonlinearly dependent on voltage, decrements in bAP amplitude may result in marked reductions in dendritic calcium channel activation with consequences for synaptic plasticity.

**Figure 2:**
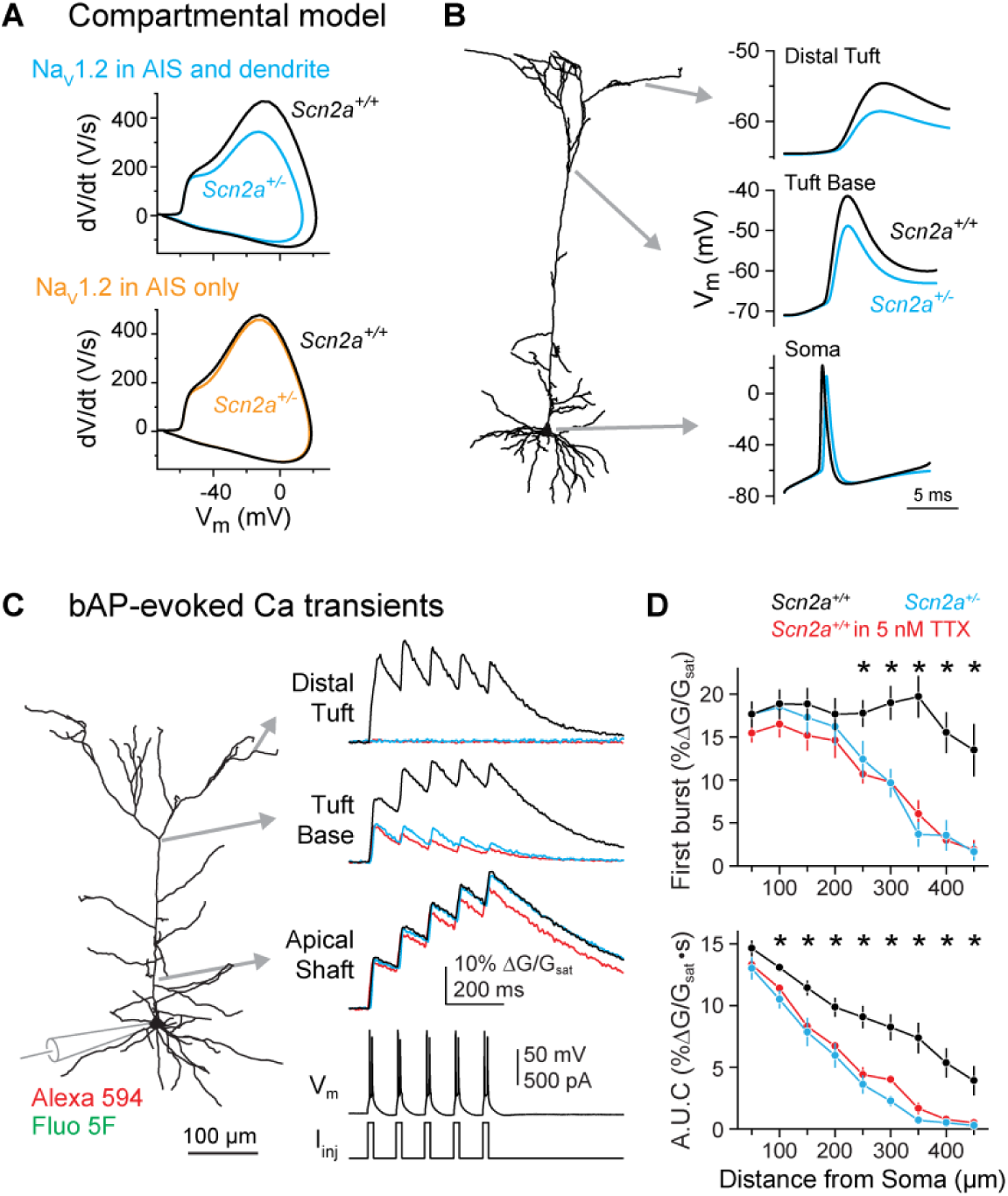
*Scn2a* haploinsufficiency impairs dendritic backpropagation of APs. **A:** Compartmental model of cortical layer 5 pyramidal cell with two distributions of NaV1.2 and NaV1.6 (see Fig. S4 for other model configurations). Top model has NaV1.2 in the proximal AIS, NaV1.6 in the distal AIS, and NaV1.2 and 1.6 equally co-expressed in the somatodendritic compartment. Note reduction in peak rising dV/dt. Bottom model has NaV1.2 in the proximal AIS only, with NaV1.6 in the somatodendritic and distal AIS compartments. Removal of half the NaV1.2 channels results in only minor changes to AP phase-plane. **B:** A single AP evoked in the top model in (A), backpropagating throughout dendrite. More marked differences in AP shape observed in more distal dendritic locations. **C:** 2PLSM calcium imaging throughout apical dendrite of L5 thick-tufted neuron. Calcium transients evoked by bursts of AP doublets. **D:** Transient amplitude is plotted for the first of 5 bursts (top) and area under the curve from stimulus onset to stimulus offset +100 ms (bottom) in *Scn2a^+/+^* (n = 10), *Scn2a^+/-^* (n = 8), and *Scn2a^+/+^* cells treated with 5 nM TTX (n = 5). Circles and bars are means ± SEM. *: p < 0.05, Kruskal-Wallis test.

To determine whether dendritic excitability was impaired in *Scn2a^+/-^* neurons, we imaged calcium transients evoked by bursts of APs at various locations throughout the apical dendrite of layer 5 pyramidal cells. In WT neurons, bursts reliably evoked calcium transients throughout the apical dendrite. By contrast, calcium transients in *Scn2a^+/-^* neurons rapidly diminished in amplitude with increasing distance from the soma, becoming virtually absent in the most distal dendrites (Fig. 2C-D). To determine whether these effects were due to acute loss of Na_V_ channels, rather than a consequence of altered excitability during development, we partially blocked sodium channels in mature WT neurons with tetrodotoxin (TTX), using a concentration that mimicked the reduced AP speed observed in *Scn2a^+/-^* neurons (5 nM, Fig. S4F-G). Consistent with acute effects of *Scn2a* haploinsufficiency, bAP-evoked calcium transients were reduced to a comparable extent in TTX-exposed WT neurons (Fig. 2C-D). Thus, *Scn2a^+/-^* haploinsufficiency leads to a major, persistent deficit in dendritic excitability in adult L5 neurons that is likely due to loss of Na_V_1.2 channels localized to the dendrite.

### Excitatory synapses are immature in *Scn2a^+/-^* neurons

Impairments of both axonal and dendritic excitability could affect synapses, which are a common locus of dysfunction in mouse models of ASD (Bourgeron, 2015; Monteiro and Feng, 2017; Tsai et al., 2012). We tested this by first measuring miniature excitatory and inhibitory postsynaptic currents (mEPSC, mIPSC) at P6 and P27, capturing developmental periods of axonal and dendritic excitability deficits, respectively. Neither mEPSCs nor mIPSCs were affected at P6. At P27, mEPSC frequency was reduced by 48% (WT: 9.3 ± 1.1 Hz, n = 23; Scn2a+/-: 4.9 ± 0.8, n = 19; p < 0.001 Mann-Whitney), with no change in mEPSC amplitude or mIPSC amplitude or frequency (Fig. 3A-B, S5A-B).

Lower mEPSC frequencies could reflect reductions in release probability or the number of functional synapses contributing to AMPA-mediated mEPSCs. The paired pulse ratio, which is sensitive to differences in release probability, was no different between *Scn2a^+/-^* and WT cells (Fig. 3C, S5C-D). By contrast, the ratio of AMPA- to NMDA-mediated current in evoked EPSCs was substantially reduced in *Scn2a^+/-^* cells (Fig. 3D; WT: 5.5 ± 0.7, n = 8: *Scn2a^+/-^*: 3.2 ± 0.2, n = 12; p < 0.01, Mann-Whitney). Taken together, the reduction in both mEPSC frequency and AMPA:NMDA ratio suggest that mature *Scn2a^+/-^* neurons have an excess of AMPA-lacking spines more commonly observed at earlier developmental time points (Kerchner and Nicoll, 2008). Consistent with these functional observations, synapse structure was also immature. While overall dendritic morphology and spine density were unaltered in *Scn2a^+/-^* neurons, spines visualized on both the apical and basal dendrites were longer and had smaller heads relative to their total head and neck volume (Fig. 4). Thus, these data indicate excitatory inputs to pyramidal cells are functionally and structurally immature in *Scn2a^+/-^* neurons.

**Figure 3:**
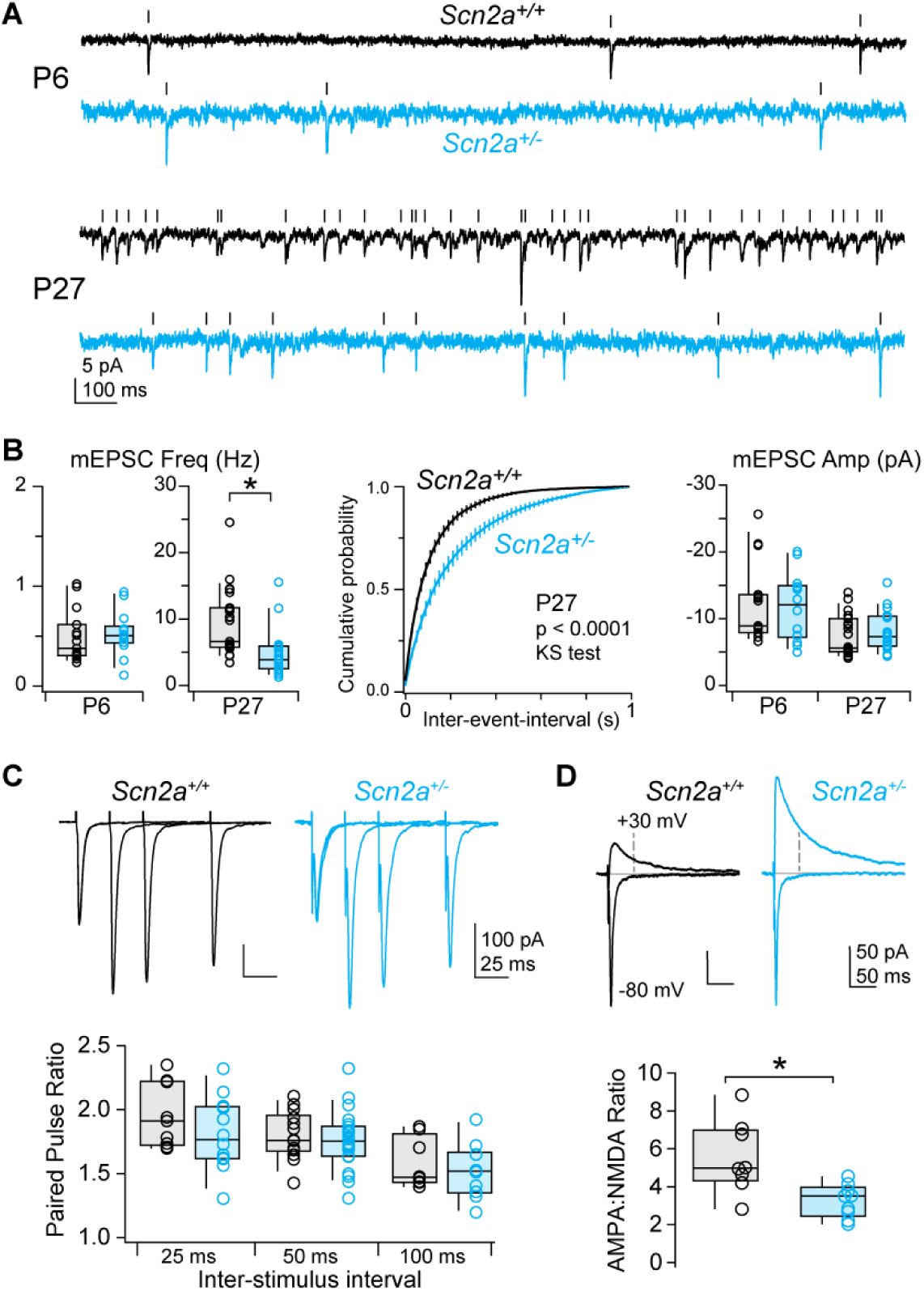
*Scn2a* haploinsufficiency disrupts excitatory synapse function. **A:** mEPSCs recorded in *Scn2a^+/+^* (black) and *Scn2a^+/-^* (cyan) pyramidal cells at P6 and P27. Tick marks denote detected events. **B:** Left, average mEPSC frequency per cell (open circles). Box plots are median, quartiles, and 90% tails. Note that frequency range is different for P6 and P27. Middle, cumulative probability distribution of mEPSC event intervals at P27. Distributions were generated per cell, then averaged. Bars are SEM. P < 0.0001, Kolmogorov-Smirnov test. Right, average mEPSC amplitude per cell. **C:** Paired pulse ratio of evoked excitatory inputs to *Scn2a^+/+^* (black) and *Scn2a^+/-^* (cyan) pyramidal cells at 3 different stimulus intervals. Bottom, summary grouped by inter-stimulus interval. No differences noted. **D:** Top, AMPA receptor-mediated and mixed AMPA/NMDA receptor-mediated evoked EPSCs at -80 and +30 mV, respectively. Dashed line denotes time at which NMDA receptor-mediated component was calculated. Bottom, AMPA-NMDA ratio in *Scn2a^+/+^* and *Scn2a^+/-^* neurons. * p < 0.01, Two-sided Mann-Whitney.

**Figure 4:**
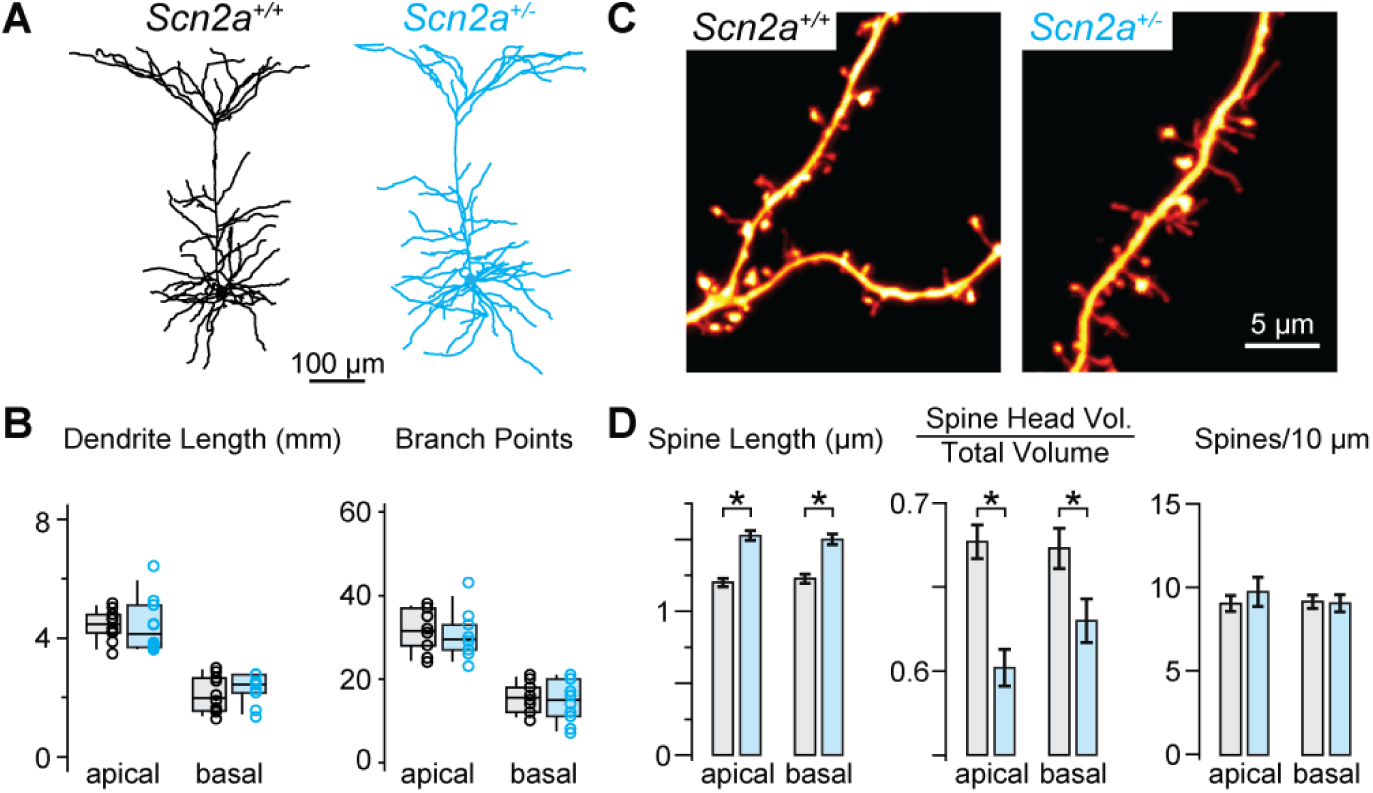
Dendritic spines are morphologically immature in *Scn2a^+/-^* cells. **A:** Examples of dendritic morphology in mature *Scn2a^+/+^* (black) and *Scn2a^+/-^* (cyan) pyramidal cells. **B:** Overall dendrite length and branch points in apical and basal trees. **C:** Examples of spines along apical dendrites. Open circles are single cells. Box plots are median, quartiles, and 90% tails. No differences across genotype noted. **D:** Overall spine length from shaft to spine head, volume of spine head relative to total volume of head and shaft, and number of spines per length of dendrite were measured in apical and basal dendrites (n = 500-600 spines per group, 1–3 dendritic branches per cell, 6 cells per group). *: p < 0.05, Two-sided Mann-Whitney.

To determine whether impaired synaptic function was due to early developmental deficits in axonal excitability or persistent deficits in dendritic excitability, we engineered a mouse with one *Scn2a* allele under Cre-loxP control (*Scn2a^+/fl^*) (Fig. S6A). Mice were first crossed to the CaMKIIα-Cre driver line, which expresses Cre in neocortical pyramidal cells only after P10 (Fig. S6B, Xu et al., 2000). As such, *Scn2a^+/fl^*::CaMKIIα-Cre mice develop with normal AP threshold and spike output. In *Scn2a^+/fl^*::CaMKIIα-Cre mice, peak dV/dt was no different at P18, likely due to a combination of low Cre expression and slow Na_V_1.2 turnover. By P50, peak dV/dt matched constitutive *Scn2a^+/-^* neurons (Fig. 5B). Furthermore, bAP-evoked dendritic Ca transients were suppressed to comparable extents as observed in constitutive *Scn2a^+/-^* mice (Fig. 5C-D). Thus, these mice developed without early deficits in axonal excitability and exhibited impaired dendritic excitability only later in development.

**Figure 5:**
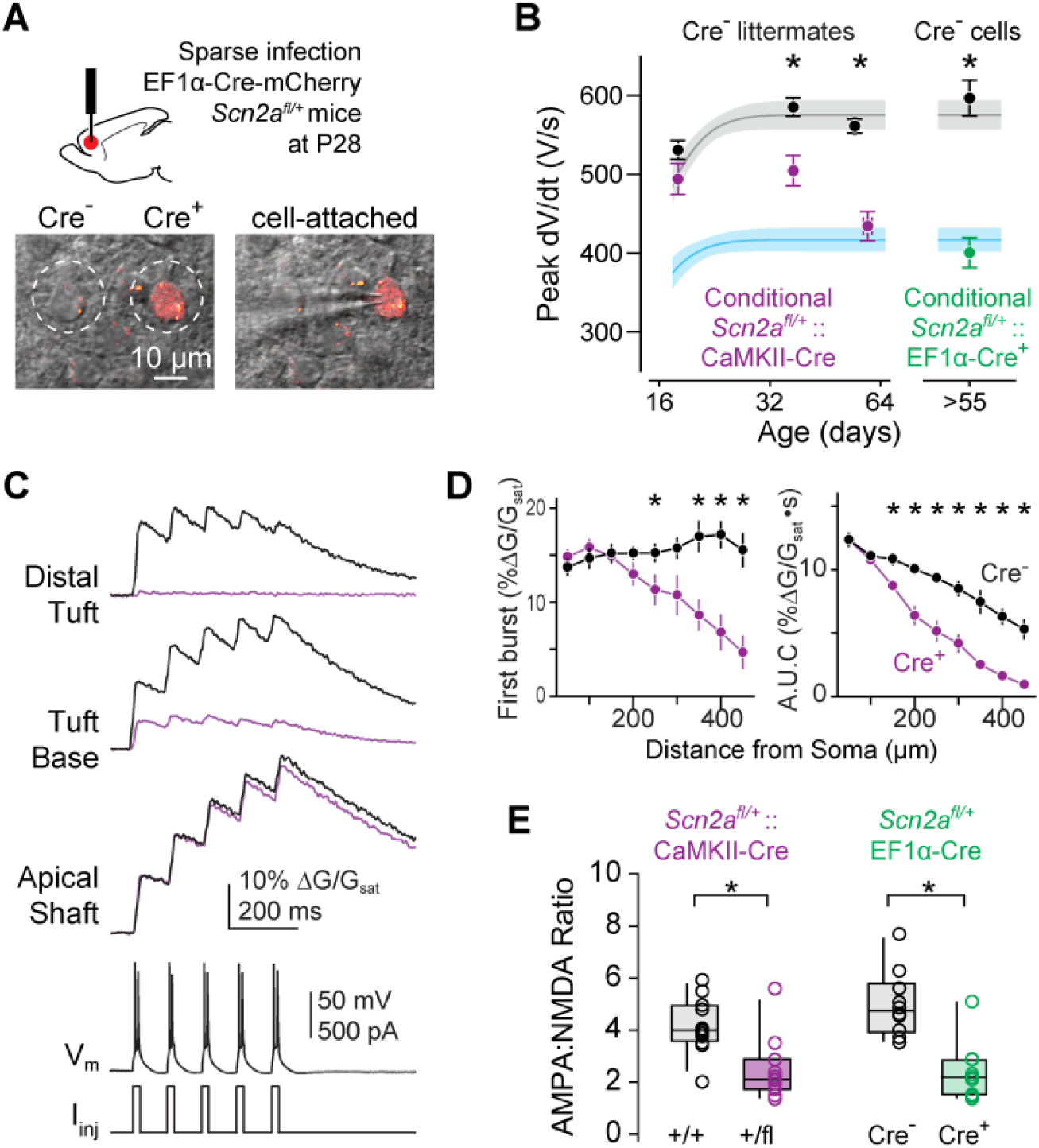
Inducing *Scn2a* haploinsufficiency late in development disrupts dendritic excitability, AP backpropagation, and synaptic stability. **A:** *Scn2a^+/fl^* mice were injected with a dilute Cre virus, infecting a subset of neurons in PFC. Images are single 2PLSM optical section of transmitted laser light (“scanning-DIC”) and mCherry fluorescence (red). Left, dashed circles highlight two neighboring Cre+ and Cre-neurons. Right, Cre+ neuron targeted for whole-cell recording with pipette from left-hand side. Image taken while in cell-attached configuration. **B:** Peak dV/dt vs age in conditional *Scn2a* mice. dVdt measurements were made from CaMKII::*Scn2a^+/fl^* or ^+/+^ mice at several ages and compared to developmental curve derived from constitutive mice. Note overlap with wild-type developmental curve at P18 and eventual overlap with *Scn2a^+/-^* curve at >P50 for CaMKII::*Scn2a^+/fl^* neurons. Neurons in *Scn2a^+/fl^* mice injected with dilute Cre virus were assessed after P55. Circles and bars are means ± SEM. *: p < 0.01, Two-sided Mann-Whitney. **C:** Dendritic calcium imaging as in Fig. 2C, except in CaMKII::*Scn2a^+/fl^* or ^+/+^ cells. **D:** Transient amplitude is plotted for the first of 5 bursts (left) and area under the curve from stimulus onset to stimulus onset + 500ms (right) in CaMKII::*Scn2a^+/+^* (n = 9 cells) or CaMKII::*Scn2a^+/-^* (n = 8 cells). Circles and bars are means ± SEM. *: p < 0.05, Two-sided Mann-Whitney. **E:** AMPA:NMDA ratio between CaMK-Cre:*Scn2a^+/+^* and *Scn2a^+/fl^* littermates, and between Cre+ and Cre-neurons in *Scn2a^+/fl^* mice injected with EF1α-Cre. * p < 0.05, Two-sided Mann-Whitney.

Strikingly, AMPA:NMDA ratio was reduced in *Scn2a^+/fl^*::CaMKIIα-Cre neurons when measured after P50, revealing that impaired dendritic excitability alone is sufficient to destabilize synapses (Fig. 5E; WT: 4.2 ± 0.3, n = 12; floxed: 2.5 ± 0.4, n = 11; p < 0.01, Two-sided Mann-Whitney). To test whether these synaptic deficits result from a cell-autonomous loss of *Scn2a*, we injected a dilute Cre-expressing adeno-associated virus (AAV-EF1α-Cre-mCherry) into mPFC of P28 *Scn2a^+/fl^* mice (Fig. 5A). Four weeks later, we found that both peak dV/dt and AMPA:NMDA ratio were reduced in Cre-positive, but not Cre-negative neurons. (Fig. 5B, E; WT: 5.0 ± 0.4, n = 10; floxed: 2.4 ± 0.4 n = 9; p < 0.01, two-sided Mann-Whitney). These results indicate that the persistent, cell-autonomous, dendritic function of Na_V_1.2 is required to maintain synaptic stability in mature pyramidal cells.

### *Scn2a^+/-^* mice have synaptic and behavioral learning deficits

Impaired synaptic stability could arise if synapses cannot be strengthened. Indeed, physiologically relevant forms of synaptic plasticity depend on the coincident detection of bAPs and glutamate at the synapse (Feldman, 2012). To test whether plasticity is impaired in *Scn2a^+/-^* cells, we induced long-term potentiation (LTP) of putative apical dendrite synapses by pairing layer 1 fiber stimulation with bursts of APs (Tzounopoulos et al., 2004). In WT neurons, paired stimulation resulted in long-lasting synaptic potentiation (Fig. 6). In contrast, LTP was not observed in either *Scn2a^+/-^* neurons or in WT neurons in 5 nM TTX (EPSP slope, normalized to baseline: WT: 1.52 ± 0.10, n = 11; *Scn2a^+/-^*: 0.98 ± 0.11, n = 11; TTX: 0.97 ± 0.06, n = 9; p = 0.0002, Kruskal-Wallis test; WT vs *Scn2a^+/-^*: p = 0.0007, WT vs TTX: p = 0.002, Dunn’s multiple comparisons test). Thus, *Scn2a* haploinsufficiency impairs synaptic plasticity, consistent with impairments in bAP-evoked calcium transients in the dendrites.

**Figure 6:**
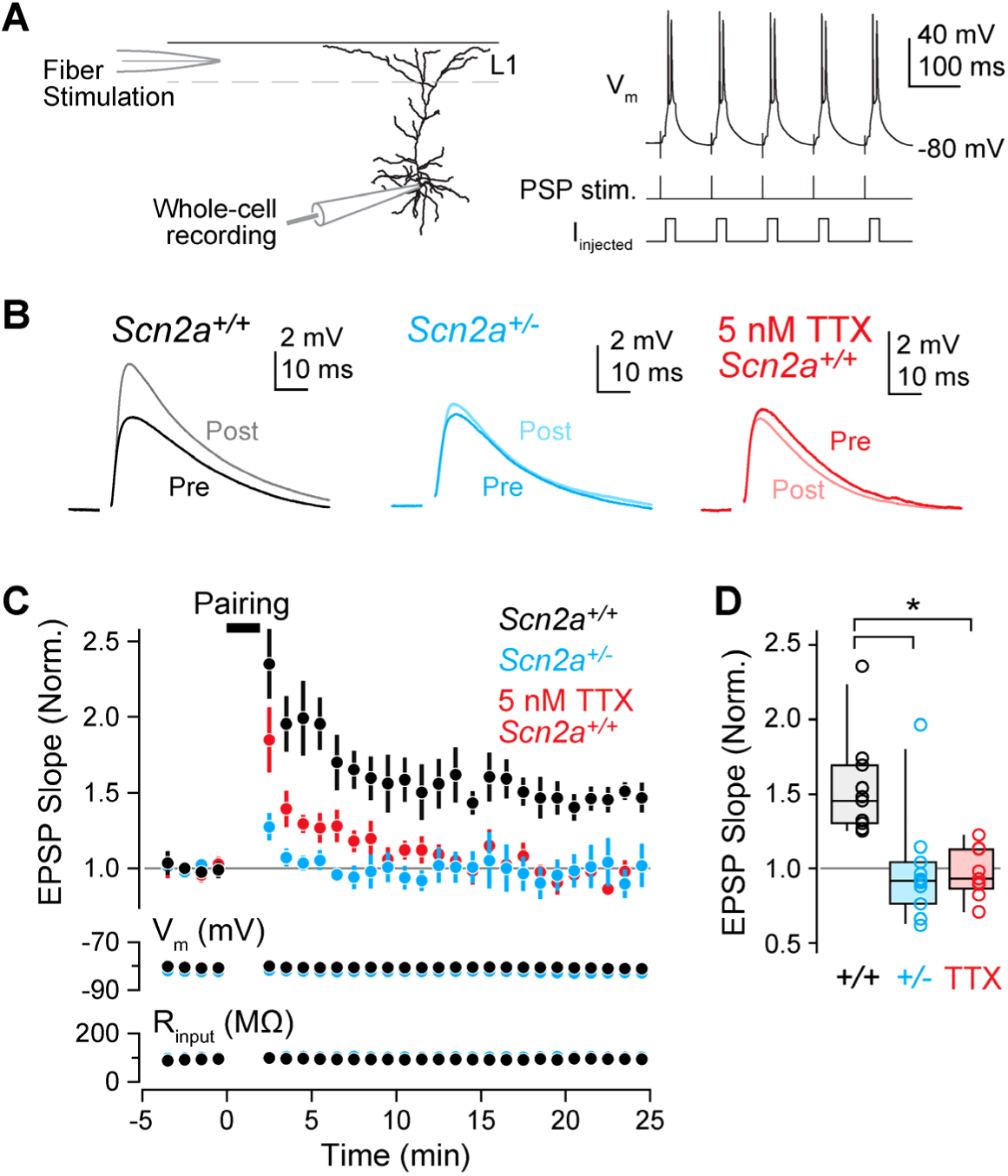
*Scn2a* haploinsufficiency impairs synaptic plasticity. **A:** Recording configuration for burst-based LTP experiments. Stimulating electrode was placed in layer 1 and synaptic stimulation was paired with somatic current-evoked AP bursts. **B:** Example of EPSPs before and after LTP pairing protocol. Lighter shades are post-induction data (15–25 minutes after pairing). Stimulus artifact removed for clarity. **C:** EPSP slope (first 2 ms), membrane potential (V_m_) and input resistance (R_in_) vs time before and after LTP induction. Circles and bars are mean ± SEM. **D:** EPSC slope per cell in *Scn2a^+/+^* cells with and without TTX and *Scn2a^+/-^* cells. *: p = 0.0002, Kruskal-Wallis test; WT vs *Scn2a^+/-^*: p = 0.0007, WT vs TTX: p = 0.002, Dunn’s multiple comparisons test.

Given these synaptic plasticity deficits, we next asked if *Scn2a* haploinsufficiency results in behavioral impairments. We assessed this by screening *Scn2a^+/-^* mice and WT littermates of both sexes through a behavioral panel designed to assess locomotion, anxiety, repetitive behavior, sociability, and learning (Fig. 7). While most behavioral assays were no different between genotypes (Fig. S7), *Scn2a^+/-^* mice exhibited sexually dimorphic impairments in reversal learning and social preference. In a water T-maze task, mice must first learn which arm of a maze contains a submerged platform (day 1–4, 4 trials/day), then reverse their association when the platform location is switched (day 5–8, 6 trials/day). Male, but not female, *Scn2a^+/-^* mice had impaired performance in the reversal phase of the task (Fig. 7A). By contrast, female, but not male, *Scn2a^+/-^* mice had social preference deficits (Fig. 7B-D). This was assessed using a two-chamber social approach task where mice had the option of interacting with a caged, sex-matched stimulus mouse in one chamber or an inanimate toy mouse in the other. Mice were paired with the same stimulus mouse for three trials, during which mice of both sexes and genotypes exhibited initial social preference with gradual habituation. When the toy was replaced with a novel stimulus mouse on the fourth trial, both male and female WT mice switched preference to the new stimulus mouse whereas female *Scn2a^+/-^* mice showed no preference for the novel mouse. These results suggest that *Scn2a^+/-^* mice have impaired behavioral flexibility and social discrimination.

**Figure 7:**
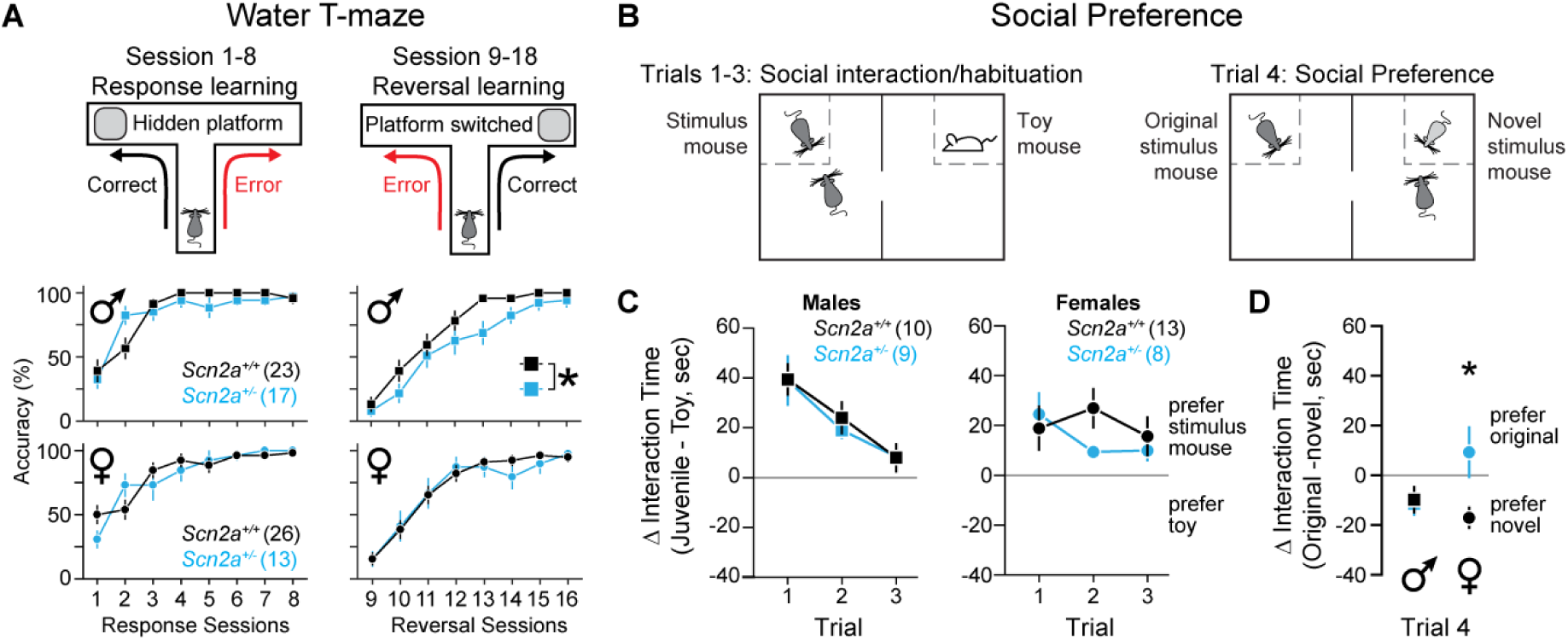
*Scn2a* haploinsufficiency impairs learning and social preference. **A:** Top: Schematic of the water T-maze behavioral assay. Mice were placed at the start of a water T-maze and had to identify the arm containing a submerged platform. Accuracy was measured as percentage of trials with correct arm choices. Bottom: Task accuracy for male and female *Scn2a^+/+^* and *Scn2a^+/-^* mice across response sessions (2 trials per session) and reversal sessions (3 trials per session). Squares/circles and bars are mean ± SEM. *: Difference between genotypes, p = 0.042, F_Genotype_ = 4.4, 2-way repeated measures ANOVA. **B:** Schematic of social preference assay. Mice were placed in a two-chamber container and time spent interacting with a stimulus mouse (same mouse each trial) and a toy mouse (trials 1–3, social interaction/habituation) or novel stimulus mouse (trial 4, social preference) was recorded. **C:** Difference in interaction time of the stimulus mouse and the toy mouse across trials 1–3 for male and female *Scn2a^+/+^* and *Scn2a^+/-^* mice. Squares/circles and bars are mean ± SEM. No statistical differences noted. **D:** Difference in interaction of the familiar stimulus mouse and a novel stimulus mouse for male and female *Scn2a^+/+^* and *Scn2a^+/-^* mice. Squares/circles and bars are mean ± SEM. *: p=0.017, Two-sided Mann-Whitney.

## Discussion

Here, we report that Na_V_1.2 channels are critical for maintaining synaptic stability through an unexpected role in dendritic excitability, and that cell-autonomous *Scn2a* haploinsufficiency late in development is sufficient to impair excitatory synaptic connectivity. This reframes *SCN2A* as a gene not only important for axonal excitability, but also as a gene essential for synaptic function, with consequences for ASD/ID etiology.

While the expression of Na_V_1.2 in the AIS is well established, there is less consensus on its distribution in pyramidal cell dendrites. The first immunostaining of Na_V_1.2 demonstrated somatic and apical dendritic localization in cortical pyramidal cells, though this may represent a pool of newly synthesized channels within vesicles (Gong et al., 1999). Subsequent investigation in hippocampal pyramidal cells found that Na_V_1.2 co-localized with the presynaptic terminal marker VGlut1 using diffraction-limited immunofluorescence techniques, and further showed that Na_V_1.6 is expressed throughout dendrites using freeze-fracture immuno-electron microscopy (Lorincz and Nusser, 2010). By contrast, more recent immuno-electron microscopy identified Na_V_1.2 throughout dendrites, with an enrichment near postsynaptic densities in spines (Johnson et al., 2017). Whether such apposition with presynaptic terminals accounts for VGlut1 co-localization is unclear. Functionally, somatodendritic Na_V_ currents are best described by Na_V_1.2 channels (Hu and Bean, 2018; Hu et al., 2009). Here, we found that AP waveform was augmented in constitutive and conditional *Scn2a^+/-^* pyramidal cells in a manner that was best explained by somatodendritic loss of Na_V_1.2, with no functional compensation from the residual *Scn2a* allele or Na_V_1.6 (Fig. 1–2, S1, S4). Moreover, AP-evoked calcium transients were suppressed throughout dendritic arbors (Fig. 2, 5). Thus, these data indicate that Na_V_1.2 is functionally expressed in the soma and dendrites of pyramidal neurons, where they play an important role in excitability and synapse function.

### Relationship with other ASD-associated genes

Though *SCN2A* has the strongest evidence of ASD association of any gene identified via exome sequencing (Ben-Shalom et al., 2017; De Rubeis et al., 2014; Sanders et al., 2015), the mechanistic underpinnings of this strong association have been a mystery. Since Na_V_1.2 channels were best understood for their role in AP initiation (Bender and Trussell, 2012; Kole and Stuart, 2012), it was difficult to find parallels with other ASD-associated genes that are associated with synaptic function and gene transcription. Data shown here instead suggest that *SCN2A* function in ASD converges in the dendrite with the large group of ASD-associated genes involved in synaptic function (De Rubeis et al., 2014). Indeed, disruptions in both excitatory synaptic function and neuronal excitability have been reported in other ASD-associated genes (Bateup et al., 2011; Bozdagi et al., 2010; Contractor et al., 2015; Meredith and Mansvelder, 2010; Nestor and Hoffman, 2012; Williams et al., 2015; Yi et al., 2016). Other ASD-associated genes play a role in regulating gene expression (De Rubeis et al., 2014). This may include the regulation of *SCN2A*. For example, interactions with *SCN2A* have been reported for both FMRP and CHD8 (Darnell et al., 2011; Sugathan et al., 2014).

*SCN2A* was one of the first genes identified with a clear excess of missense mutations linked to ASD and ID (Ben-Shalom et al., 2017; Sanders et al., 2015). Recent efforts have identified other genes harboring ASD-associated missense variants, though the functional implications of such mutations are largely unclear (Geisheker et al., 2017). Na_V_1.2 function, by contrast, can be well characterized, allowing for a better understanding of how differences in channel activity affect neuronal function. Several recurring *SCN2A* missense variants block sodium flux (Ben-Shalom et al., 2017). These may be functionally identical to protein truncating variants, since we and others find that protein levels are reduced 50% in *Scn2a^+/-^* mice (Planells-Cases et al., 2000), with no evidence for functional compensation. Other missense variants observed in ASD patients alter channel voltage dependence or kinetics, reducing neuronal excitability in models of developing cortical pyramidal cells (Ben-Shalom et al., 2017). Whether they result in similar dendritic deficits in mature neurons remains to be tested.

### Role of Scn2a in synapse function and learning

*Scn2a* haploinsufficiency selectively impairs dendritic excitability and synaptic function in mature PFC pyramidal cells, with no effects on AP initiation or propagation (Fig. 1, S2). This distinguishes Na_V_1.2 from other Na_V_ isoforms expressed in the mature cortex that either support overall excitability in inhibitory neurons (Na_V_1.1) or contribute to both action potential generation and dendritic excitability in pyramidal neurons (Na_V_1.6) (Hu et al., 2009; Lorincz and Nusser, 2008; Ogiwara et al., 2007). Therefore, in the mature brain, Na_V_1.2 may play a major role in supporting the backpropagation of action potentials within pyramidal cell dendrites. Here, we demonstrated that Na_V_1.2 channels are critical for supporting bAPs from the AIS to the distal dendrites. In addition, Na_V_1.2 may also promote the forward propagation of synaptic potentials from synapses to the AIS. Dendritic sodium channels have been reported to amplify the amplitude of EPSPs recorded at the soma and to contribute to the generation of dendritic sodium spikes that can occur in the absence of somatic APs (Araya et al., 2007; Golding and Spruston, 1998; Lipowsky et al., 1996; Sun et al., 2014). The degree to which Na_V_1.2 supports these functions is unknown, but its disruption may further compound issues in ASD/ID.

Given the ubiquity of *Scn2a* expression in pyramidal neurons in both the neocortex and hippocampus, it is likely that the consequences of *Scn2a* haploinsufficiency extend to regions beyond PFC. Indeed, cortical visual impairments are often comorbid in children with loss-of-function *SCN2A* variants (Sanders et al., 2018), perhaps due to deficits in sensory cortex. Furthermore, *Scn2a^+/-^* mice exhibit truncated hippocampal replay events during sharp-wave ripples and have related deficits in spatial learning (Middleton et al., 2018). While cellular mechanisms underlying these effects remain unexplored, impaired dendritic excitability and synaptic connectivity could contribute to both the altered replay and spatial learning deficit.

Deficits in behavioral flexibility and social recall identified here were sexually dimorphic, appearing in males and females, respectively. Reversal learning deficits we observed in male mice parallel spatial learning impairments observed previously in male *Scn2a^+/-^* mice (Middleton et al., 2018). Whether female *Scn2a^+/-^* are similarly resistant to spatial learning impairments has not been explored. Nevertheless, these data add to a growing body of literature highlighting sex-based differences in the behavior of neurodevelopmental disorder models (Angelakos et al., 2017; Dhamne et al., 2017; Grissom et al., 2018; Reith et al., 2013; Schmeisser et al., 2012; Zamarbide et al., 2018), paralleling the sex biases observed in ASD and, to a lesser extent, ID (Fischbach and Lord, 2010; Kim et al., 2011). While we provide insight into how cortical physiology is altered by *Scn2a* loss, future work will be needed to elucidate the cellular and synaptic consequences of *Scn2a* haploinsufficiency in regions outside of PFC, and whether region-specific impairments underlie behavioral deficits in a sex-specific manner.

In conclusion, we found that haploinsufficiency induction at two different developmental timepoints, both well after Na_V_1.2 channels cease to contribute to AP initiation, was sufficient to alter synaptic function (Fig. 3, 5). While this result does not exclude additional effects on network function due to reduced axonal excitability in early cortical development, it does indicate that persistent Na_V_1.2 function in maintaining dendritic excitability is critical for proper synaptic strength. Thus, restoring proper Na_V_1.2 function, even in the mature brain, may be an enticing avenue for therapeutic intervention in loss-of-function *SCN2A* cases.

## Acknowledgments

We are grateful to Drs. E. Glasscock and M. Montal for providing *Scn2a^+/-^* mice, and to Drs. G. Davis, K. Kay, D. Manoli, M. Scanziani and members of the Bender and Sanders labs for critically assessing this work. Behavioral data were obtained with the help of the Gladstone Institutes’ Neurobehavioral Core. CaMKII-Cre::Ai14 image acquisition and analysis of dendritic spines was performed at the Gladstone Institutes’ Histology & Light Microscopy Core.

### Funding

This research was supported by SFARI grant 513133 (KJB), the Natural Sciences and Engineering Research Council (NSERC) of Canada PGS-D Scholarship (PWES), National Institutes of Health Grant No. F32 NS095580 (RBS) and R01 MH110928 (SJS).

### Author contributions

Conceptualization: PWES, RBS, SMS, KJB; Methodology: PWES, RBS, KJB; Software: PWES, RBS, KJB Jr, RLC; Formal Analysis: PWES, RBS, CMK, KJB; Investigation: PWES, RBS, CMK, KJB Jr, KJB; Resources: SMS, KJB; Writing—original draft: PWES, KJB; Writing—review & editing: all authors; Visualization: PWES, RBS, KJB; Supervision: KJB; Project Administration: SMS, KJB; Funding Acquisition: PWES, RBS, SMS, KJB.

## Supplemental Figures

**Supplemental Figure 1:**
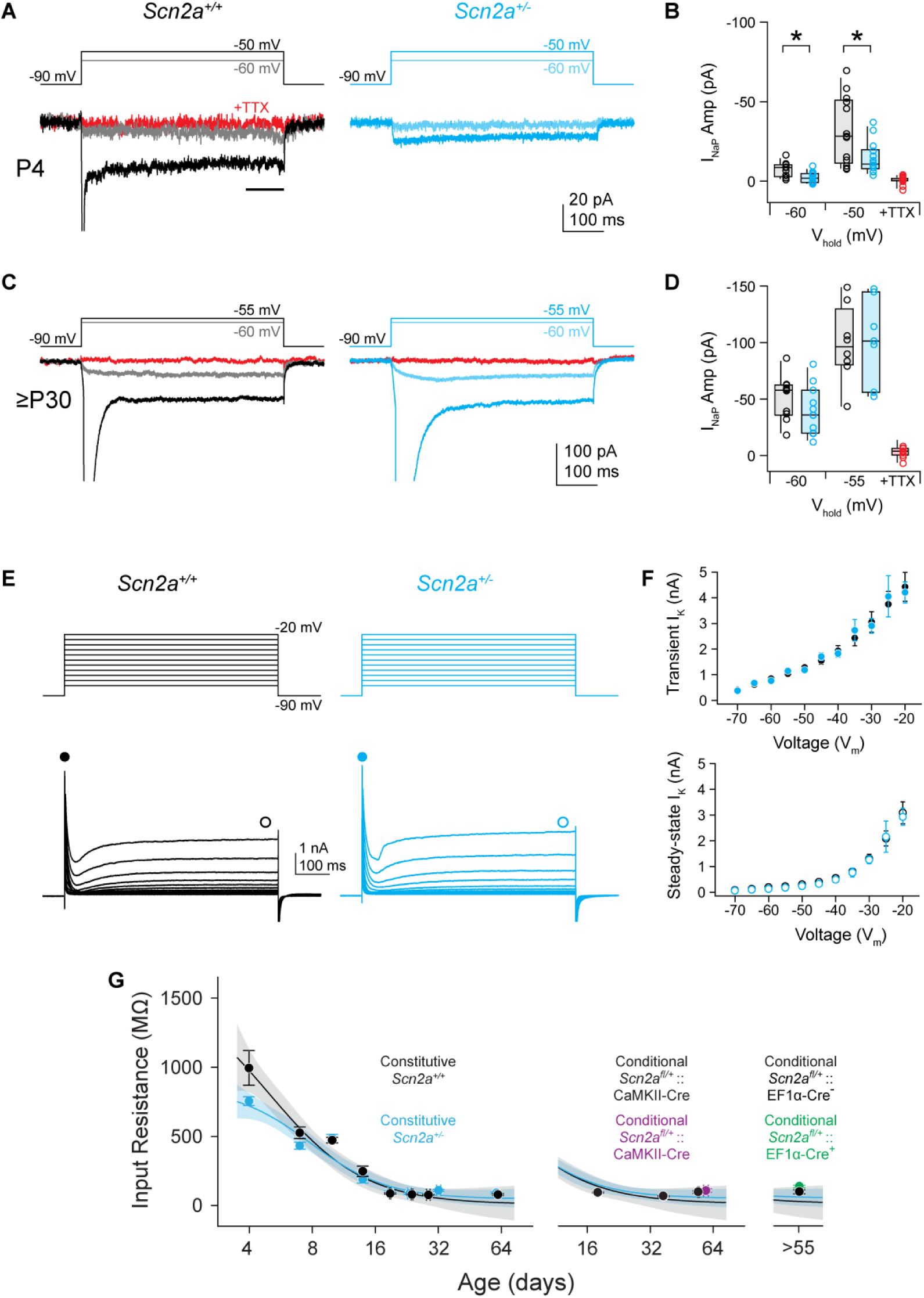
Additional electrophysiological measurements in *Scn2a^+/-^* neurons. **A:** At P4, NaV-mediated currents were evoked with voltage steps from -90 mV to either -60 or -50 mV. Note generation of transient current at -50 mV. Data are color coded to match voltage step. Currents generated in TTX were assessed with steps from -90 to -50 mV only. Horizontal black bar highlights timing of persistent currents measurement. **B:** Summary of persistent current amplitudes. Open circles are single cells. Box plots are median, quartiles, and 90% tails. TTX was applied in a subset of cells, and data were pooled over both *Scn2a^+/+^* and *Scn2a^+/-^* cells. *: p < 0.05, Two-sided Mann-Whitney. **C:** NaV-mediated currents from P30-40 neurons. Note that voltage steps range from -90 to either -60 or -55, instead of -50, due to increased current amplitude and voltage-dependent differences between NaV1.2 presumably recruited in P4 neurons and NaV1.6 presumably recruited in P3-40 neurons. **D:** Summary as in *b*, but for P30-40 neurons. No differences observed between *Scn2a^+/+^* and *Scn2a^+/-^* cells at this age range. **E:** Potassium currents generated from voltage steps from -90 to -20 mV (10 mV increments) in *Scn2a^+/+^* and *Scn2a^+/-^* cells. Closed and open circles denote when transient and steady-state current amplitudes were analyzed. Data obtained from P32-39 animals. **F:** Summary of current amplitudes per voltage step. Circles and bars are mean ± SEM. No differences noted across genotype. **G:** Left: Development of input resistance (AP max-threshold) in *Scn2a^+/+^* and *Scn2a^+/-^* cells. Data fit with sigmoid function with 95% confidence bands based on group mean values. Circles and bars are mean ± SEM within an age group. Middle: development of action potential amplitude of *Scn2a^fl/+^*::CaMKII-cre and WT mice. Right: Action potential amplitude of Cre+ and Cre-*Scn2a^fl/+^* neurons sparsely infected with AAV-EF1α-Cre-mCherry at P28. No statistical differences noted.

**Supplemental Figure 2:**
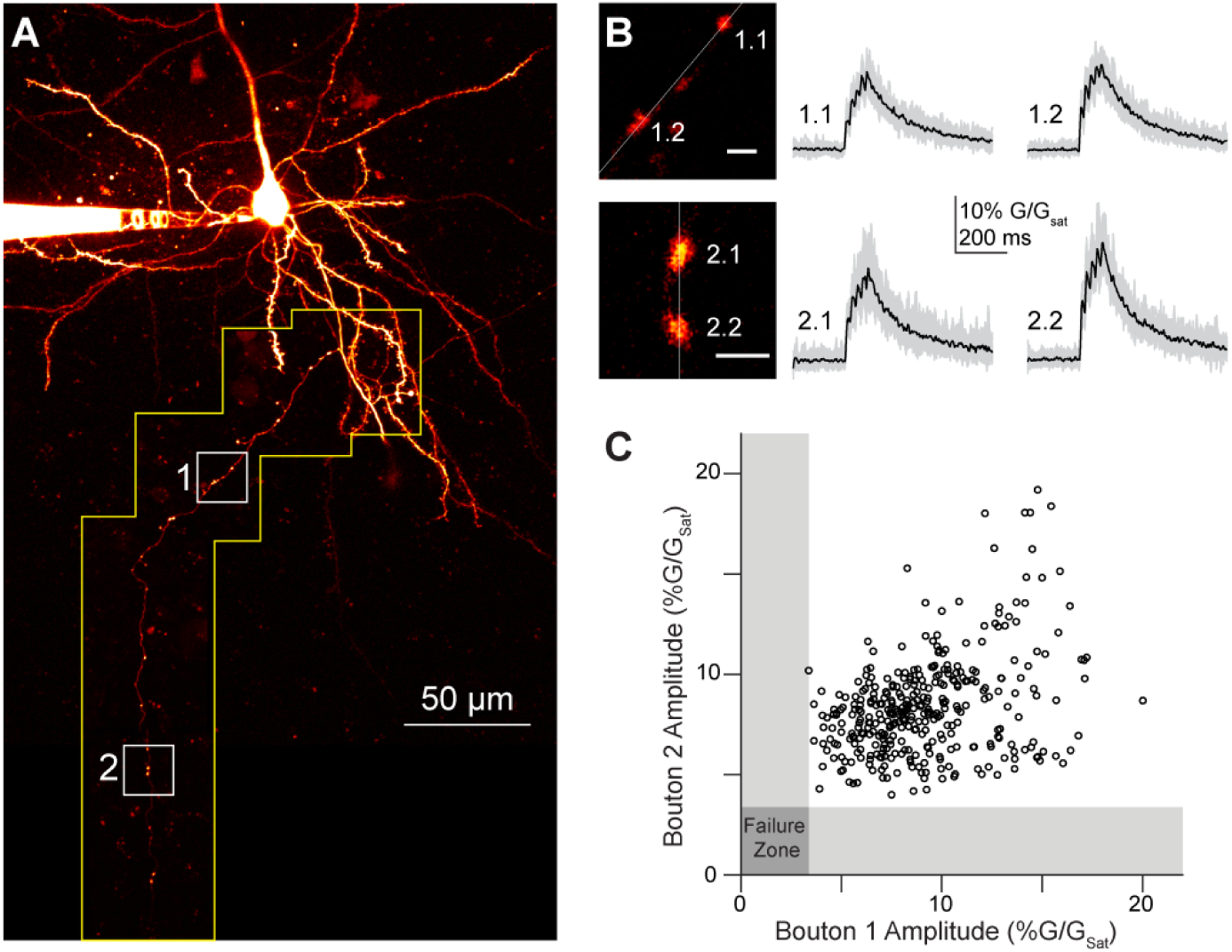
APs reliably evoke calcium transients in axonal boutons of *Scn2a^+/-^* pyramidal cells. **A:** 2PLSM z-stack of basal dendrites and axon of a mature pyramidal cell filled with alexa 594. Inset: bouton pairs used for calcium imaging **B:** Calcium transients of boutons pairs highlighted in A in response to a train of 5 action potentials at 20Hz. Grey traces are individual sweeps, black are averages of 20 sweeps. **C:** Comparison of calcium transient amplitudes of individual sweeps for each bouton pair in response to a single action potential (e.g., the first of the 5 APs shown in B). The coincident failure the spike-evoked calcium transient to exceed 2 standard deviations above the baseline signal (highlighted by the grey box) was considered an action potential failure. No coincident failures were observed. 15-20 trials per bouton pairs, 4–8 bouton pairs per cell, 3 cells.

**Supplemental Figure 3:**
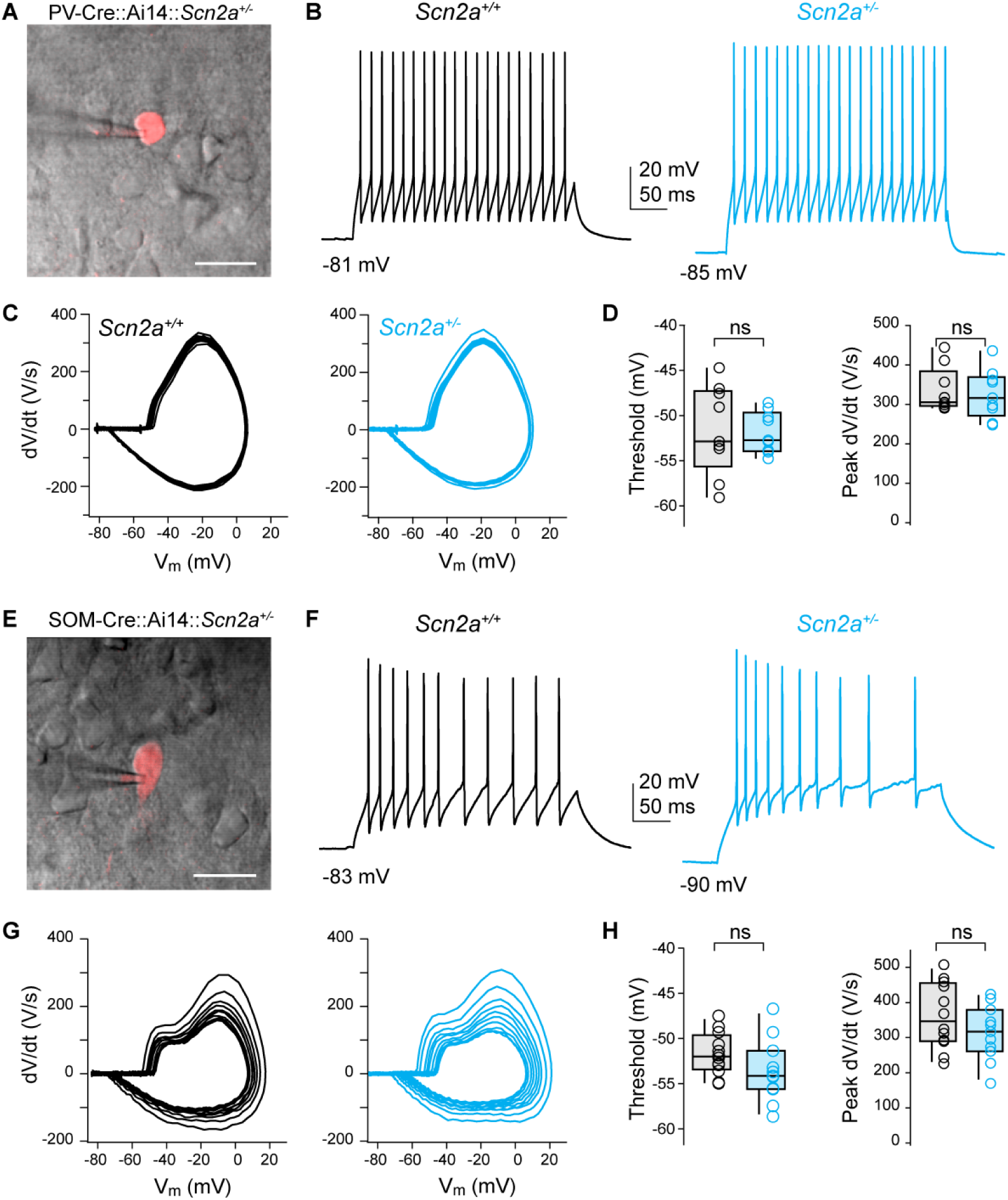
Layer 5 parvalbumin and somatostatin neuron APs are not altered in *Scn2a^+/-^* mice. **A:** 2PLSM single optical section of tdTomato-positive parvalbumin interneuron, overlaid with scanning-DIC image showing pipette in cell-attached configuration. Scale bar: 20 µm. **B:** Spiking generated from parvalbumin positive cells in *Scn2a^+/+^* and *Scn2a^+/-^* mice. **C:** Phase-plane plots of data shown in (b). **D:** AP threshold and peak rising dV/dt for the first AP of spike train. Open circles are single cells. Box plots are median, quartiles, and 90% tails. No statistical differences noted. Data obtained from P34-40 animals. **E-H:** Identical to (a-d), but for somatostatin positive interneurons. Data obtained from P37-38 animals.

**Supplemental Figure 4:**
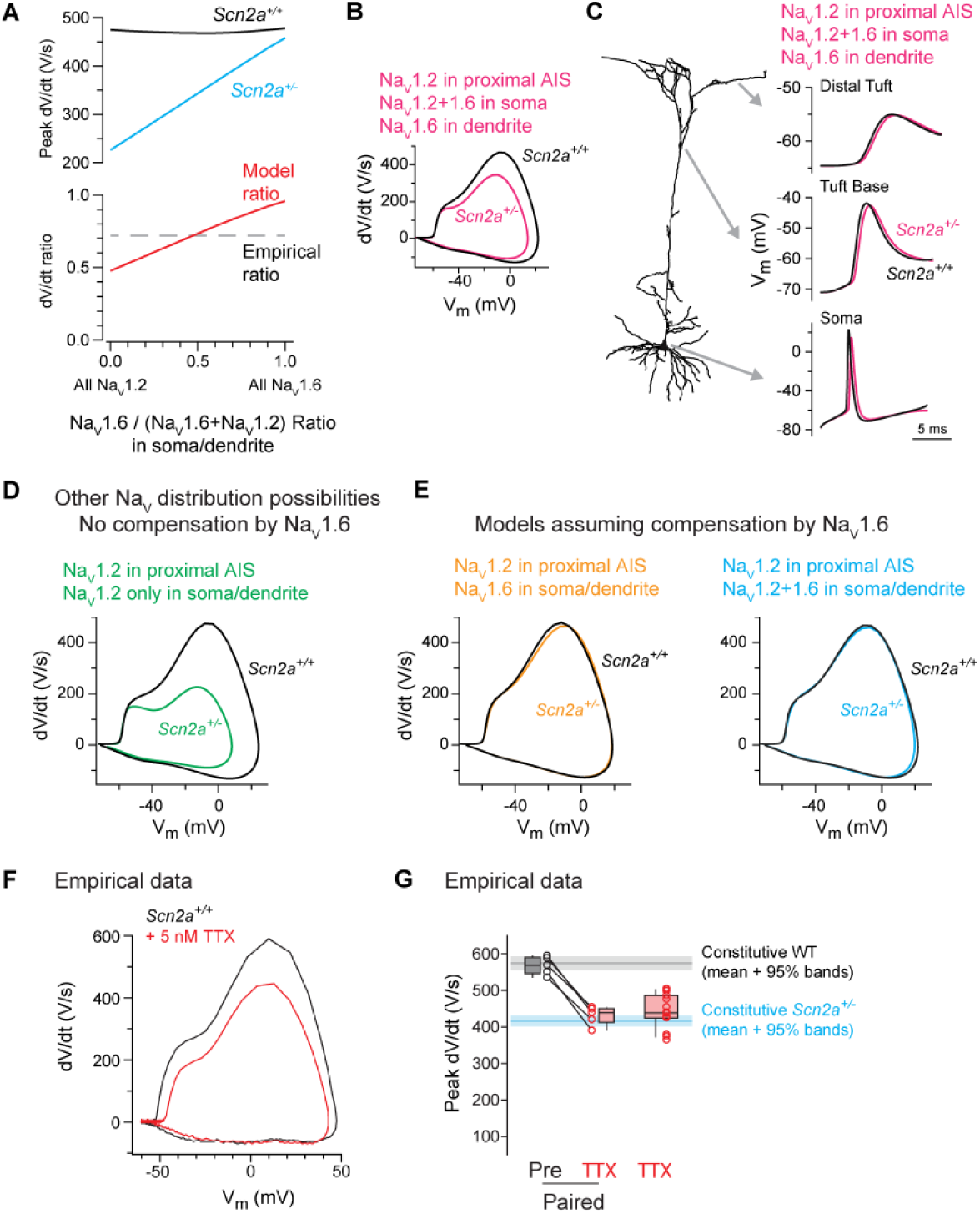
Modeling NaV1.2 loss in pyramidal cells with different NaV compartmental distributions and mimicking *Scn2a^+/-^* with sub-saturating TTX. **A:** Top, the peak rising phase dV/dt in cells expressing different ratios of NaV1.2 vs NaV1.6 in the somatodendritic compartment in WT and *Scn2a^+/-^* conditions. Bottom, ratio of *Scn2a^+/-^* to WT peak dV/dt vs. ratio of NaV1.2 vs NaV1.6 in the somatodendritic compartment. Larger relative reductions in peak dV/dt are observed in cases where NaV1.2 dominates the somatodendritic NaV conductance. The ratio that best matches empirical observations (dashed grey line) occurs when equal numbers of NaV1.2 and NaV1.6 are expressed in the somatodendritic compartment. **B:** Compartmental model of cortical layer 5 pyramidal cell where NaV1.2 expression is restricted to the proximal AIS and is co-expressed in equal levels with NaV1.6 in the soma. In this model configuration, NaV1.6 is the only channel isoform in the dendrite. Note that both models resemble empirical observations, suggesting that proximal AIS and somatic NaV1.2 channel distributions largely influence phase plane shape recorded at the soma. **C:** Backpropagation of single spikes for conditions modeled in (B). AP waveform is shown at several points along dendritic arbor, including the base of the apical tuft and the distal part of the apical tuft. No major differences are observed in backpropagation when NaV1.2 is only in the AIS and soma (right). Thus, while this model can account for empirically observed differences in somatically recorded dV/dt, it cannot account for deficits in bAP evoked dendritic Ca transients (Fig. 2). **D:** Left, Model in which NaV1.2 is the only NaV isoform expressed throughout the somatodendritic compartment. *Scn2a* haploinsufficiency in this condition alters dV/dt far more dramatically than observed experimentally. **E:** Models in which NaV1.2 loss in different compartments is compensated by increased expression of NaV1.6. Two cases modeled in Fig. 2 are considered (same color code). Note marginal change to phase plane plot, suggesting that compensation is unlikely in *Scn2a^+/-^* conditions. **F:** Example phase-plane plots of action potential from a cell pre and post treatment with 5nM TTX. **G:** Peak action potential velocity for cells pre and post 5nM TTX and cells recorded with 5nM TTX in recording solution at the time of whole-cell recording initiation. Data compared to mean and 95% confidence intervals of *Scn2a^+/+^* (black/grey) and *Scn2a^+/-^* (blue) neurons (Fig. 1F).

**Supplemental Figure 5:**
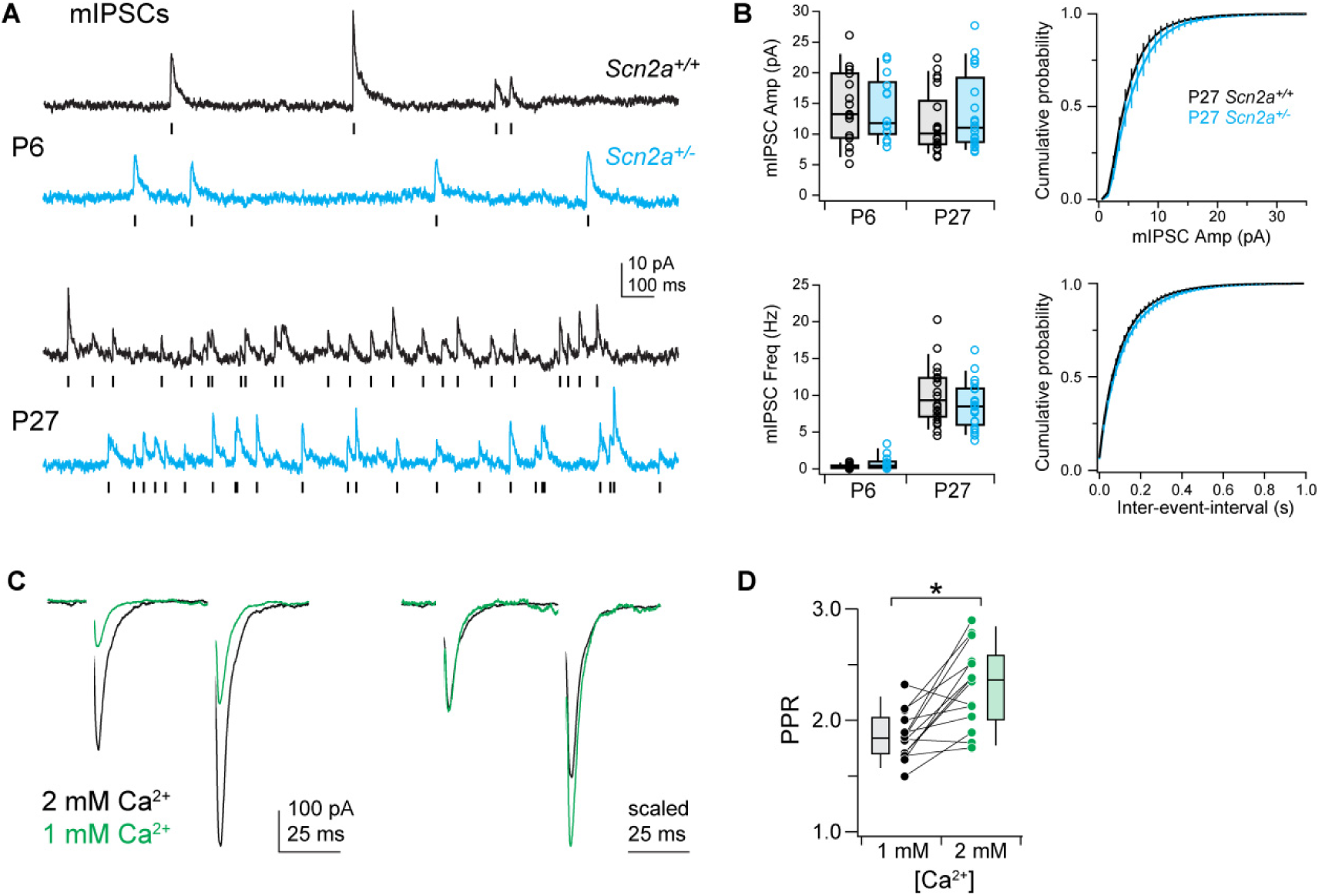
mIPSCs and PPR controls. **A:** Examples of mIPSCs recorded in pyramidal cells voltage-clamped to 0 mV in *Scn2a^+/+^* (black) and *Scn2a^+/-^* (cyan) cells. Tick marks denote detected events. Data obtained at P6 (top) and P27 (bottom). **B:** Mean mIPSC frequency and amplitude per cell, separated by age. Open circles are single cells. Box plots are median, quartiles, and 90% tails. Cumulative probability histograms were generated per cell, then averaged across all cells. Bars are SEM per data bin. n = 15, 14, 22, 21 for P6 *Scn2a^+/+^*, P6 *Scn2a^+/-^*, P27 *Scn2a^+/+^,* and P27 *Scn2a^+/-^* cells, respectively. No statistical differences noted. **C:** Reducing extracellular calcium from 2 to 1 mM (divalents compensated with MgCl2) in WT neurons reduces EPSC amplitude an increases PPR, demonstrating PPR is sensitive to changes in release probability at these synapses. Left, raw traces. Right, scaled to highlight increase in PPR. Stimulus artifact blanked for clarity. **D:** Summary of PPR change evoked by altering extracellular calcium. Circles are single cells. Lines connect data obtained from single cells. Box plots are median, quartiles, and 90% tails. Data obtained at 50 ms ISI only. Asterisk: p = 0.004, Wilcoxon signed rank test, n = 12 cells.

**Supplemental Figure 6:**
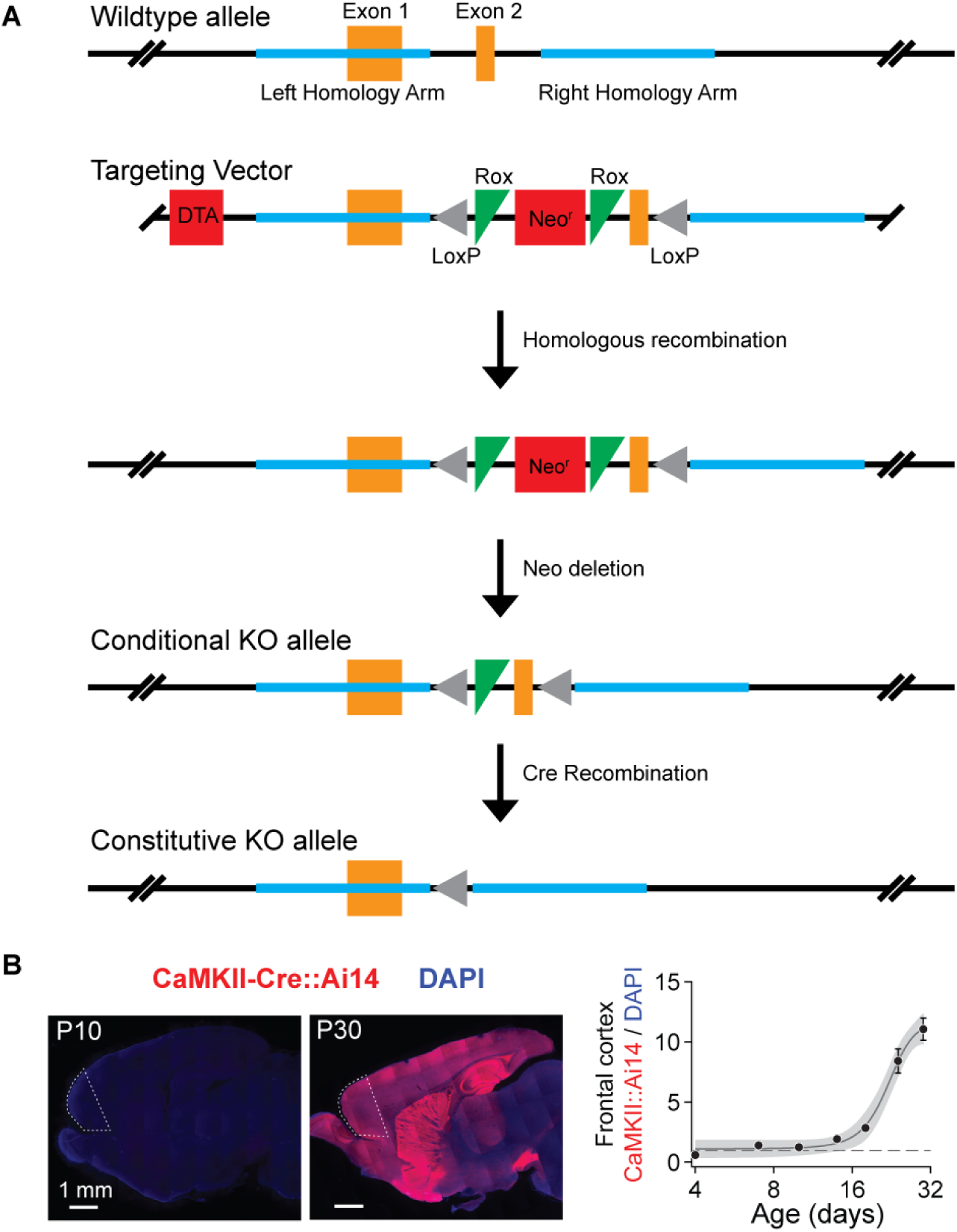
Conditional Mouse Construction. **A:** Schematic overviewing the construction of the conditional *Scn2a^+/fl^* allele. Briefly, LoxP sites were inserted around exon 2 of the Scn2a gene by homologous recombination in embryonic stem (ES) cells and screened for successful insertion. Positive ES cells were introduced into host embryos and transferred to surrogate mothers, generating F0 chimeras that were outbred with C57BL/6 to generate heterozygous mutants. Please see methods for further details. **B:** Left, CaMKII-Cre mice were crossed to Ai14 reporter mice, and tdTomato-mediated fluorescence was measured across development from P4-P32 to determine when Cre-driven tdTomato expression can first be observed in frontal cortex of CaMKII-Cre mice. Values were normalized to DAPI intensity. Measurements were made from frontal cortex regions encapsulated by dashed white perimeter. Right, Ai14/DAPI fluorescence ratio over development, normalized to P4 data. Circles and bars are mean ± SEM at each age (n = 6 sections from 2 animals). Data fit with sigmoid function with 95 confidence bands.

**Supplemental Figure 7:**
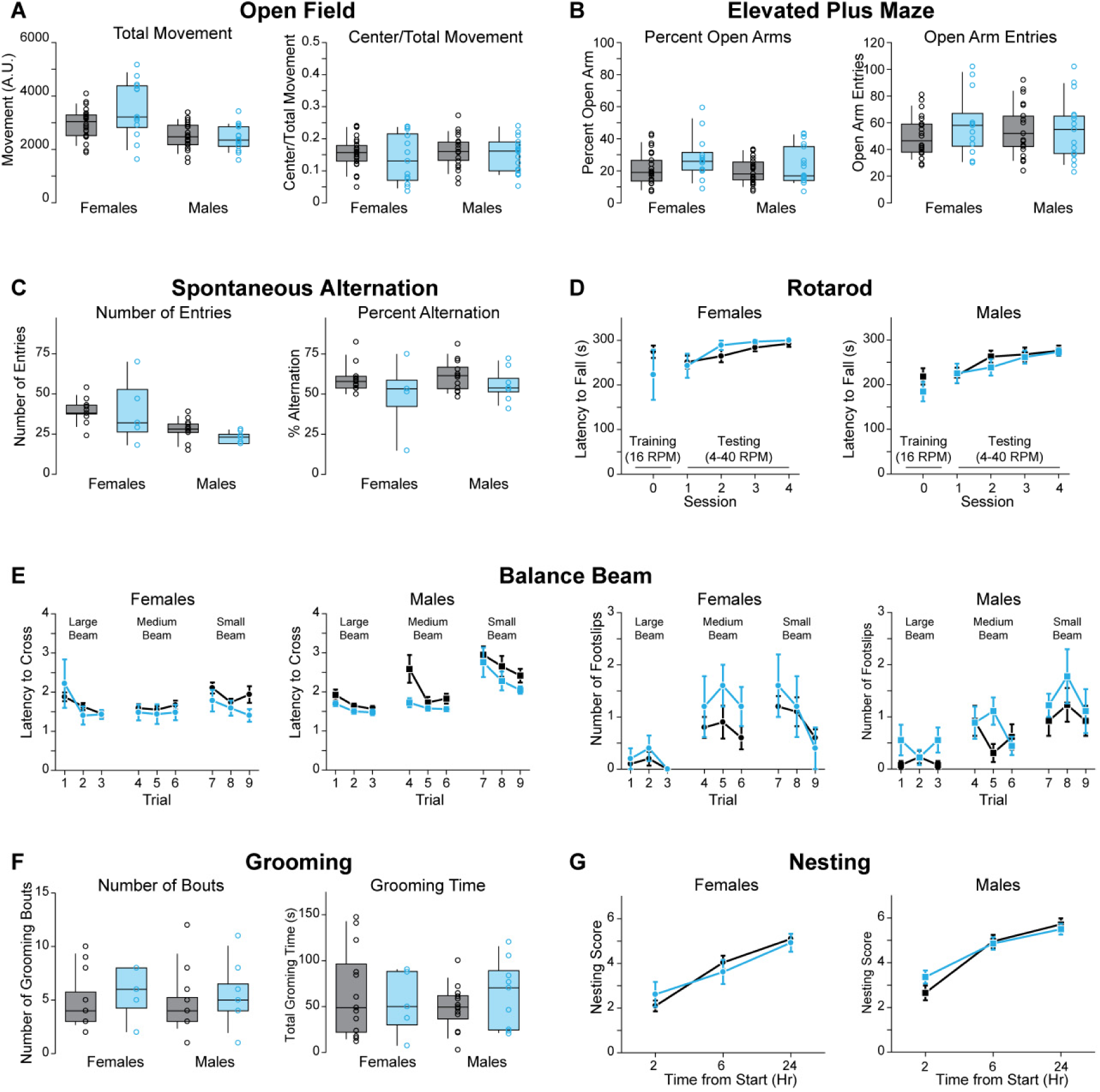
*Scn2a^+/-^* mice have normal ambulation, anxiety, spontaneous alternation, motor coordination, balance, grooming, and nesting behavior. **A:** Left: total movement and Right: proportion of movement in the center during an open field test. Comparisons are of male and female *Scn2a^+/+^* (23 male, 26 female) and *Scn2a^+/-^* (18 male, 13 female) mice. **B:** Left: total time spent in the open arms and Right: total number of open arm entries during an elevated plus maze test. Comparisons are of male and female *Scn2a^+/+^* (23 male, 26 female) and *Scn2a^+/-^* (18 male, 13 female) mice. **C:** Left: number of arm entries made and Right: percent of arm entries contributing to entering all three arms of a Y-maze in succession without repeating an arm (total number of alternation/(total number of entries – 2) *100). Comparisons are of male and female *Scn2a^+/+^* (13 males, 13 females) and *Scn2a^+/-^* (9 males, 5 females) mice. **D:** Latency to fall during rotarod training (16 RPM) and testing (variable speed, 4–40RPM). Comparisons are of male and female *Scn2a^+/+^* (13 males, 10 females) and *Scn2a^+/-^* (9 males, 5 females) mice. **F:** Left: latency to cross and Right: number of foot slips while crossing a small, medium or large balance beam (three trials per beam). Comparisons are of male and female *Scn2a^+/+^* (13 males, 13 females) and *Scn2a^+/-^* (9 males, 5 females) mice. **G:** Left: number of grooming bouts and Right: total grooming time of isolated mice over a 10-minute undisturbed period. Comparisons are of male and female *Scn2a^+/+^* (13 males, 13 females) and *Scn2a^+/-^* (9 males, 5 females) mice. **G:** Nesting scores of singly housed mice measured at 2, 6, and 24 hours from the start of nest creation. Comparisons are of male and female *Scn2a^+/+^* (23 males, 26 females) and *Scn2a^+/-^* (18 males, 13 females) mice. No statistical differences noted.

## Materials and methods

All experimental procedures were performed in accordance with UCSF and Gladstone Institutes IACUC guidelines. *Scn2a^+/fl^* mice were created by inserting LoxP sites around exon 2 of the *Scn2a* gene (Cyagen). A targeting vector was generated by PCR using BAC clone RP23-332C13 and RP24-42717 from the C57BL/6J library to create homology arms around a LoxP flanked cKO region that includes exon 2 of the *Scn2a* gene. The targeting vector included a Neo cassette flanked by Rox sites for positive selection, and DTA outside of the homology arms for negative selection. The linearized vector was subsequently delivered to ES cells (C57BL/6) via electroporation, followed by drug selection, PCR screening, and Southern Blot confirmation. Confirmed clones were then introduced into host embryos and transferred to surrogate mothers. Chimerism in the resulting pups was identified via coat color. F0 male chimeras were bred with C57BL/6 females to generate F1 heterozygous mutants that were identified by PCR.

For slice electrophysiology and 2-photon imaging, mice aged P4 through P62 were anesthetized, and 250 µm-thick coronal slices containing prefrontal cortex were prepared. Slices were prepared from *Scn2a*^+/-^, CaMKII-Cre (MGI: 2446639)::*Scn2a^+/fl^*, Parv-Cre (MGI: 3590684)::Ai14::*Scn2a^+/-^*, SOM-Cre (MGI: 4838416)::Ai14::*Scn2a^+/-^*, or *Scn2a* wild type littermates (genotyped by PCR). All data were acquired and analyzed blind to *Scn2a* genotype, except for experiments examining the effects of tetrodotoxin in a wild type background (AP waveform and synaptic plasticity), and the effects of sparse Cre transfection in *Scn2a^+/fl^* mice. Data were acquired from both sexes (blind to sex), with no sex-dependent differences noted in measurements made in acute slice recordings [e.g., AMPA:NMDA ratio, combined data across constitutive and conditional heterozygote cases: Male +/+ 5.1 ± 0.3, n = 8, Female +/+: 4.7 ± 0.3, n = 22, Male +/-: 2.4 ± 0.3, = 18, Female +/-: 3.1 ± 0.3, n = 11; p = 0.13, 2-factor ANOVA. Peak rising dV/dt of AP, combined data from P23-62 constitutive and >P50 conditional heterozygote cases: Male +/+: 584.4 ± 8.8 V/s, n = 48; Female +/+: 588.0 ± 22.8, n = 6; Male +/-: 405.3 ± 8.1, n = 32; Female +/-: 449 ± 19.0, n = 9; p = 0.21, 2-factor ANOVA). Cutting solution contained (in mM): 87 NaCl, 25 NaHCO_3_, 25 glucose, 75 sucrose, 2.5 KCl, 1.25 NaH_2_PO_4_, 0.5 CaCl_2_ and 7 MgCl_2_; bubbled with 5%CO_2_/95%O_2_; 4°C. Following cutting, slices were either incubated in the same solution or in the recording solution for 30 min at 33°C, then at room temperature until recording. Recording solution contained (in mM): 125 NaCl, 2.5 KCl, 2 CaCl_2_, 1 MgCl_2_, 25 NaHCO_3_, 1.25 NaH_2_PO_4_, 25 glucose; bubbled with 5%CO_2_/95%O_2_; 32–34°C, ~310 mOsm.

Neurons were visualized with differential interference contrast (DIC) optics for conventional visually guided whole-cell recording, or with 2-photon-guided imaging of reporter-driven tdTomato fluorescence overlaid on an image of the slice (scanning DIC). For current-clamp recordings and voltage-clamp recordings of K^+^ currents, patch electrodes (Schott 8250 glass, 3–4 MΩ tip resistance) were filled with a solution containing (in mM): 113 K-Gluconate, 9 HEPES, 4.5 MgCl_2_, 0.1 EGTA, 14 Tris_2_-phosphocreatine, 4 Na_2_-ATP, 0.3 tris-GTP; ~290 mOsm, pH: 7.2–7.25. For Ca^2+^ imaging, EGTA was replaced with 250 µM Fluo-5F and 20 µM Alexa 594. For voltage-clamp recordings of persistent Na+ currents and synaptic activity, internal solution contained (in mM): 110 CsMeSO_3_, 40 HEPES, 1 KCl, 4 NaCl, 4 Mg-ATP, 10 Na-phosphocreatine, 0.4 Na_2_-GTP, 0.1 EGTA; ~290 mOsm, pH: 7.22. All data were corrected for measured junction potentials of 12 and 11 mV in K- and Cs-based internals, respectively.

Electrophysiological data were acquired using Multiclamp 700A or 700B amplifiers (Molecular Devices) via custom routines in IgorPro (Wavemetrics). For measurements of action potential waveform, data were acquired at 50 kHz and filtered at 20 kHz. For all other measurements, data were acquired at 10–20 kHz and filtered at 3–10 kHz. For current-clamp recordings, pipette capacitance was compensated by 50% of the fast capacitance measured under gigaohm seal conditions in voltage-clamp prior to establishing a whole-cell configuration, and the bridge was balanced. For voltage-clamp recordings, pipette capacitance was compensated completely, and series resistance was compensated 50%. Series resistance was <15 MΩ in all recordings. Experiments were omitted if input resistance changed by > ±15%.

Between P4 and P10, whole-cell current-clamp recordings were made from in the center of the cortical plate, corresponding to developing layer 5 (Cánovas et al., 2015). Putative pyramidal cells were identified based on regular spiking characteristics. Pyramidal cell identity was validated by 2-photon visualization of dendritic spines in a subset of these recordings (25/81). After P13, laminae became more distinct and recordings were restricted to L5b. In pyramidal cells aged >P17, AP characteristics are known to vary based cell class (Clarkson et al., 2017). To minimize variability, recordings were restricted to cells with low or high HCN expression levels, corresponding to intratelencephalic (IT) or pyramidal tract (PT) neurons, respectively. In current clamp, PT neurons were defined as those that exhibited a voltage rebound more depolarized that V_rest_ following a strong hyperpolarizing current (−400 pA, 120 ms) that peaked within 90 ms of current offset (*42*). All others were defined as IT. These metrics were not employed in cells from <P15 due to a lack of mature HCN-mediated current. AP threshold and peak dV/dt measurements were determined from the first AP evoked by a near-rheobase current in pyramidal cells (300 ms duration; 10 pA increments), or the first AP within a train of APs with a minimum inter-AP frequency of 25 Hz in inhibitory neurons. Threshold was defined as the V_m_ when dV/dt measurements first exceeded 15 V/s.

Miniature excitatory and inhibitory postsynaptic currents (mEPSC, mIPSC) were assessed in the presence of 10 µM R-CPP and 400 nM TTX. mEPSCs and mIPSCs were measured while voltage-clamping neurons at -80 and 0 mV, respectively. Events were detected using a deconvolution algorithm based on Pernía-Andrade et al. (2012), with a noise threshold of 3.5x. All events larger than ±2 pA that were separated by 1 ms were analyzed and subsequently screened manually. Baseline recording noise was not different across genotype (P27 mEPSC RMS noise, WT: 2.1 ± 0.2 pA, n = 23; *Scn2a^+/-^*: 2.2 ± 0.2 pA, n = 19, p = 0.6, Two-sided Mann-Whitney test).

In experiments measuring paired pulse ratio and AMPA:NMDA ratio, EPSCs were evoked via a bipolar glass theta electrode placed ~200 µm lateral to the recorded neuron in layer 5. Paired pulse ratio was assessed in the presence of 10 µM R-CPP and 25 µM picrotoxin at -80 mV. Overall divalent concentration was maintained when calcium was lowered from 2 to 1 mM by increasing MgCl_2_ from 1 to 2 mM. For AMPA:NMDA ratio, R-CPP was omitted and neurons were held at -80 and +30 mV to assess AMPA and NMDA-mediated components. AMPA and NMDA components were defined as the peak inward current at -80 mV and the outward current 50 ms after stimulus onset at +30 mV, respectively.

For sparse Cre expression experiments, mice were anesthetized with isoflurane and positioned in a stereotaxic apparatus. 500 nL volumes of AAV-EF1A-Cre-IRES-mCherry (UNC Vector Core) diluted with saline (1:10) were injected into the mPFC of *Scn2a^fl/+^* mice (stereotaxic coordinates [mm]: anterior-posterior [AP], +1.7, mediolateral [ML] −0.35; dorsoventral [DV]: −2.6). Mice were used in experiments four weeks post injection.

Persistent Na^+^ and currents were activated with 500 ms voltage steps from −90 mV and corrected using p/n leak subtraction. 10–15 trials were averaged per voltage step. Current amplitudes were calculated as the average of the last 100 ms of each step. Experiments were performed in 25 μM picrotoxin, 10 μM NBQX, 10 mM TEA, 2 mM 4-AP, 200 μM Cd^2+^, 2 μM TTA-P2, and 1 mM Cs^+^. K^+^ currents were activated with 500 ms voltage steps from -90 to -20, in 10 mV increments. 5 trials were averaged per voltage step. Current amplitudes were calculated from the transient peak and sustained components (last 50 ms). Experiments were performed in 500 nM TTX, 25 μM picrotoxin, 10 μM NBQX, and 1 mM Cs^+^. Ca` ^2+^ channels were not blocked to allow for activation of Ca^2+^-dependent K^+^ channels.

In burst-based spike timing-dependent plasticity protocols, excitatory postsynaptic potentials (EPSPs) were evoked with a theta stimulating electrode placed in layer 1, 25–50 µm from the L1/2 border, 350–500 µm dorsal of the recorded L5b pyramidal cell. After establishing a stable baseline (EPSP ISI: 0.1 Hz), PSPs were paired with AP bursts evoked by 500–800 pA somatic current steps (20 ms, onset: 10 ms after EPSP stimulation). These EPSP-AP pairings were delivered in a train of 5 at 100 ms ISI. Trains were repeated every 5s for 20 trials. Following induction, EPSP stimulation frequency was reset to 0.1 Hz, and changes in EPSP slope were assessed by comparing data 15–25 min following induction to baseline. For experiments performed in 5 nM TTX, bursts were evoked with somatic current injection steps of larger amplitude (800–1500 pA).

Two-photon laser scanning microscopy (2PLSM) was performed as previously described (*44*). A 2-photon source (Coherent Ultra II) was tuned to 810 nm for morphology and calcium imaging. Epi- and transfluorescence signals were captured either through a 40×, 0.8 NA objective for calcium imaging or a 60×, 1.0 NA objective for spine morphology imaging, paired with a 1.4 NA oil immersion condenser (Olympus). Fluorescence was split into red and green channels using dichroic mirrors and band-pass filters (575 DCXR, ET525/70m-2p, ET620/60m-2p, Chroma). Green fluorescence (Fluo-5F) was captured with 10770–40 photomultiplier tubes selected for high quantum efficiency and low dark counts (PMTs, Hamamatsu). Red fluorescence (Alexa 594) was captured with R9110 PMTs. Data were collected in linescan mode (2–2.4 ms/line, including mirror flyback). For calcium imaging, data were presented as averages of 10–20 events per site, and expressed as 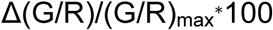, where (G/R)_max_ was the maximal fluorescence in saturating Ca^2+^ (2 mM) (Yasuda et al., 2004). Backpropogation experiments were performed in 25 μM picrotoxin, 10 μM NBQX and 10 µM R-CPP.

Spine morphology and density images were obtained during 2PLSM images obtained at 2x the Nyquist resolution limit for 810 nm excitation, with z-stacks through the dendritic arbor (0.1 µm steps in the z-axis). Stacks were post-processed with the CANDLE denoising protocol (Coupé et al., 2012), then reconstructed using IMARIS 6 (Bitplane). Maximum intensity image projections are displayed within the main and supplemental figures using the “Red Hot” lookup table (FIJI). Full dendritic reconstructions were stitched together using pairwise stitching in FIJI before generation of maximum intensity projection.

### Behavioral analyses

Open Field – Mice were transferred to the testing room 60 minutes prior to the start of testing. Mice were then placed in a clear plastic chamber (41 × 41 × 30 cm) and allowed to explore. Movement in the center and outer periphery were recorded by an array of 16 × 16 photobeams (San Diego Instruments).

Elevated plus maze – Mice were placed at the center of an elevated plus maze consisting of two open arms (5.715 cm wide, 70.485 cm long), and two enclosed arms (5.715 cm wide, 70.485 cm long, 16.51cm walls) elevated 63 cm above the ground and were allowed to explore for 10 minutes. Location, distance travelled, and arm entries were measured by infrared photobeam breaks.

Balance beam – On day 1, mice were first trained to talk across a wide beam (14.5” long x 5/8” diameter circular beam) over two guided trials. Following training, mice completed three trials (15-minute inter-trial interval) where they were placed end of the beam and traversed the wide beam unguided into an opposing dark chamber. On day 2 the mice completed three trials using a medium beam (14.5” long x 1/2” diameter circular beam), and on day 3 three trials were completed on the small beam (14.5” long x 1/4” wide square beam). For each tested trial, the latency to traverse, number of foot slips, and falls were recorded. All testing was performed under normal light conditions.

Rotarod – Mice were trained on a rotarod apparatus over three trials (inter-trial interval 15 minutes) with the rod rotating at a constant speed of 16 rotations per minute (rpm). Following training, mice were tested on 2 consecutive days with 2 sessions of 3 trials each (2-hour inter-session interval, 15-minute inter-trial interval) in which mice were placed on a rotatrod apparatus that gradually accelerated from 4 rpm to 40 rpm in 4 rpm increments every 30 seconds. Trials would end when the mouse would fall off the apparatus or after 5 minutes. All testing was performed under normal light conditions.

Nesting – Mice were placed in a standard mouse cage with 2 cm of paper chip bedding and a single nestlet (5 cm square of pressed cotton batting). Nests in the cage were then scored 2, 6, and 24 hours after the introduction of the mouse on the following scale: 0 = nestlet untouched, 1 = less than 10% of the nestlet is shredded, 2 = 10-50% of the nestlet is shredded but there is no shape to the nest, 3 = 10-50% of the nestlet is shredded and there is shape to the nest, 4 = 50-90% of the nestlet is shredded but there is no shape to the nest, 5 = 50-90% of the nestlet is shredded and there is shape to the nest, 6 = Over 90% of the nestlet is shredded but the nest is flat, 7 = Over 90% of the nestlet is shredded and the nest has walls that are as tall as the mouse on at least 50% of its sides. Half scores were given to nests that lacked well-defined walls but had clear indentations in the middle where the mouse could sit.

Spontaneous Alternation – Mice were placed in an arm of a Y-maze consisting of three arms (30 × 5.5 × 15 cm) and allowed to explore for 6 minutes. Successful alterations were counted when the mouse entered all three arms in succession without repeating an arm. Percent alternation was calculated by the total number of alterations in 6 minutes/(total number of entries - 2) * 100. Entries were defined as all four paws entering an arm.

Grooming – Mice were placed in a clear plastic container and left undisturbed for 20 minutes. Following 10 minutes of habituation, the number of grooming bouts and the total time spent grooming during the subsequent 10-minute period were then manually scored.

Response and reversal learning – Mice were singly housed and habituated to the testing room for two days prior to the start of testing. Under red light conditions, mice were placed in the start of a water T-maze (10 cm wide, 31 cm long and 17 cm tall) and were required to locate a submerged escape platform at the end of either the right of left arm of the T-maze. Mice performed 4 trials per day in 2 sessions of 2 trials each (3 hour inter-session interval and 15 minute inter-trial interval) for four consecutive days. The platform location remained in the same location across trials for each mouse, and platform location was counterbalanced between mice. On day 5, the platform location was moved to the opposite arm and mice were required to learn the new platform location. Mice performed 6 trials a day in 2 separate sessions of 3 trials (3 hour inter-session interval and 15 minute inter-trial interval) for 4 consecutive days. The maximum length of each trial was 60 seconds, and if mice failed to find the platform within the time limit they were guided to the platform. After successfully finding the platform, the mice were allowed to remain for 10 seconds. Trials were considered incorrect if the mouse entered failed to enter the correct arm first or if the trial time limit expired. All trials were recorded and tracked using Ethovision (Noldus).

Four Trial Social Preference – Testing was conducted with a white acrylic box divided into two chambers with clear acrylic dividers containing arch-shaped entrances at their center (24in. long, divided into two 12 in chambers, 16 in. wide, and 8.75 in. tall). The chambers contained 10cm square, open-bottomed social enclosures made of clear acrylic with two staggered rows of small access holes (0.5cm in diameter) 2.5cm from the bottom on the 2 sides of the enclosures facing toward the center of the chamber. Sex-matched stimulus mice were habituated to the enclosures for 3 10-minute sessions, between which they were returned to their home cages for at least 10 minutes. Both the experimental and stimulus mice were brought into testing room and given one hour to acclimate to the normal lighting conditions and room prior to testing. For trial 1, the stimulus and toy mouse were placed in the enclosures, after which the experimental mouse was placed in the chamber and given 10 minutes to freely explore the box and interact with the social enclosures. The stimulus and experimental mouse were then returned to their home cages for 1 hour before starting trial 2. Trial 2 and 3 proceeded identically to trial 1, using the same stimulus mouse in the same social enclosure. Trial 4 proceeded as in trials 1–3, however the toy mouse was replaced with a novel sex-matched stimulus mouse. Each trial was recorded by video and analyzed with Ethovision (Noldus). Interaction time was considered the total time spent sniffing within 2 cm of the social enclosures.

### Modeling

A pyramidal cell compartmental model was implemented in the NEURON environment, with baseline distributions of Na_V_1.2 and Na_V_1.6 set as previously described (Ben-Shalom et al., 2017). For phase plane comparisons, the first AP evoked with 2.2 nA stimulus intensity (100 ms duration) were compared in each model configuration. For backpropagation into dendritic arbors, stimuli were shorted to 20 ms to match empirical stimulus conditions (STDP). Stimulus intensity was increased to 2.7 nA. Relative distributions of Na_V_1.2 and Na_V_1.6 in the proximal AIS, soma and dendrite were adjusted as described in figure legends.

### Chemicals

Fluo-5F pentapotassium salt, and Alexa Fluor 594 hydrazide Na^+^ salt were from Invitrogen. Picrotoxin, R-CPP, and NBQX were from Tocris. TTX-citrate was from Alomone. All others were from Sigma.

### Statistics

Data are summarized either as single points with error bars (mean ± standard error) or with box plots depicting the median, quartiles, and 90% tails with individual datapoints overlaid. N denotes cells for all electrophysiology, spines for spine morphology, and animals for behavior. Unless specifically noted, no assumptions were made about the underlying distributions of the data and rank-based nonparametric tests were used. Statistical tests are noted throughout text. Significance was set at an alpha value of 0.05. Statistical analysis was performed using Prism (Graphpad Software), Statview (SAS), and custom routines in Matlab (Mathworks) and R.

